# Long-read only assembly of *Drechmeria coniospora* genomes reveals widespread chromosome plasticity and illustrates the limitations of current nanopore methods

**DOI:** 10.1101/866020

**Authors:** Damien Courtine, Jan Provaznik, Jerome Reboul, Guillaume Blanc, Vladimir Benes, Jonathan J. Ewbank

## Abstract

Long read sequencing is increasingly being used to determine eukaryotic genomes. We used nanopore technology to generate chromosome-level assemblies for 3 different strains of *Drechmeria coniospora*, a nematophagous fungus used extensively in the study of innate immunity in *Caenorhabditis elegans*. One natural geographical isolate demonstrated high stability over decades, whereas a second isolate, not only had a profoundly altered genome structure, but exhibited extensive instability. We conducted an in-depth analysis of sequence errors within the 3 genomes and established that even with state-of-the-art tools, nanopore methods alone are insufficient to generate eukaryotic genome sequences of sufficient accuracy to merit inclusion in public databases.

## Background

*Drechmeria coniospora* is an obligate parasitic fungus belonging to the order of Hypocreales. This fungus forms spores that adhere to the cuticle of a range of different nematodes to infect them [1]. We adopted *D*. *coniospora* strain ATCC-96282, derived from a strain isolated in Sweden, as a model pathogen for *Caenorhabditis elegans* 20 years ago [2]. We have cultured this strain, referred to here as Swe1, continuously since then, using it to understand innate immune mechanisms in its nematode host [3,4].

As part of our characterization of the interaction between *D*. *coniospora* and *C. elegans*, in 2013, we extracted DNA from our laboratory strain of the time (referred to here as Swe2), and determined its genome. Despite attempts to complete the assembly, the Swe2 genome remained fragmented, with an N50 of 3.86 Mb [5]. In addition to the genome of Swe2, a second *D*. *coniospora* genome is available (referred to here as Dan2) [6], derived from a strain related to a Danish isolate (Dan1; Figure 1). Although corresponding to a chromosome level assembly, this latter genome still contains large stretches (up to 500 kb) of undetermined sequence. In this study, we used Oxford Nanopore Technology (ONT) long-read sequencing to assemble complete fungal genomes. This revealed that the 2 isolates (Swe1 and Dan1) display strikingly different levels of genomic stability. We provide a detailed analysis that illustrates the continuing challenges to using only ONT long-read sequencing for genome assembly. As the genome sequences were of insufficient quality to allow accurate gene prediction, we polished the genomes using short DNA reads to generate high-quality sequences, providing a resource for future comparative studies.

**Figure 1.**
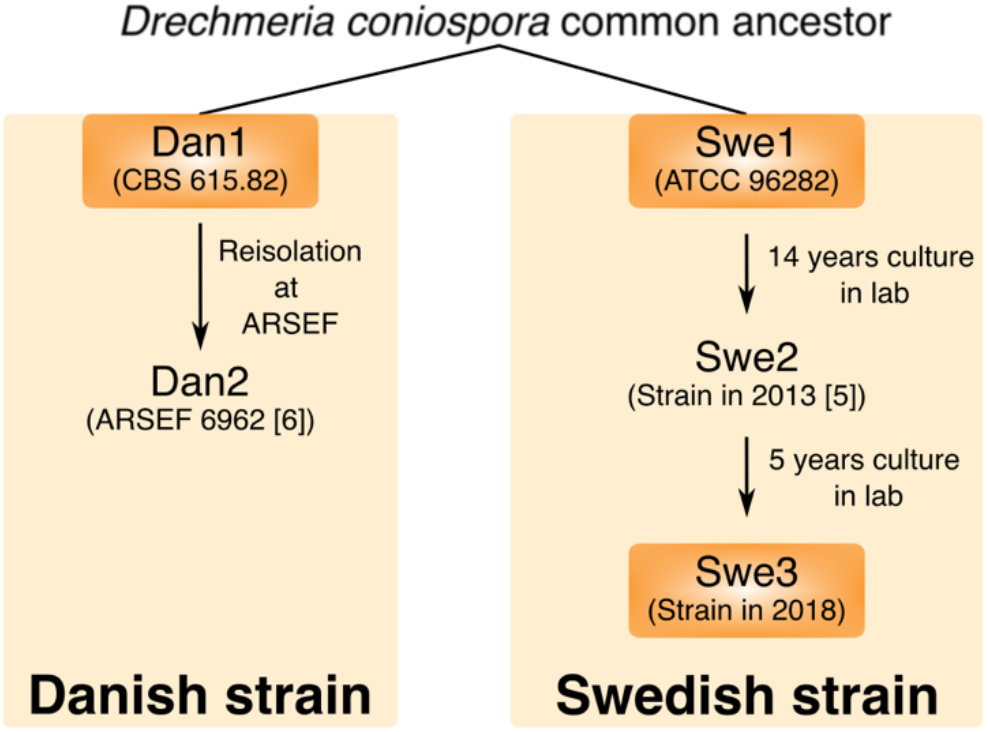
An overview of *D. coniospora* strain isolation and culture history. A strain of *D. coniospora* collected from Denmark in 1982 at the latest was deposited at the CBS-KNAW culture collection, now held by the Westerdijk Fungal Biodiversity Institute as CBS615.82. It was transferred in 1987 to the ARS Collection of Entomopathogenic Fungal Cultures (as ARSEF 2468) and then re-isolated in 2001 as ARSEF 6962. A second strain collected from Sweden was deposited at the American Type Culture Collection as ATCC 96282. It has been cultured through serial passage in *C. elegans* continuously since 1999.

## Result

An all-against-all *in silico* genome comparison of the 2 publicly available *D. coniospora* genome sequences, for Dan2 [6] and Swe2 [5], indicated the presence of extensive genomic rearrangements (Figure 2A). These could reflect real differences or assembly errors in one or both genomes. We directly confirmed one major rearrangement by PCR (Figure 2B, C), suggesting that the differences could be real. To characterise this genomic plasticity, we determined the genomes of 3 strains related to the 2 that had been sequenced previously (Figure 1). We used ONT nanopore sequencing to generate long reads and current assembly tools to construct chromosome level assemblies for all 3 strains (Supplementary Fig. S1, Supplementary Table S1). Manual curation allowed complete ca. 30 kb mitochondrial genomes to be predicted from the assemblies generated by Canu [7].

**Figure 2.**
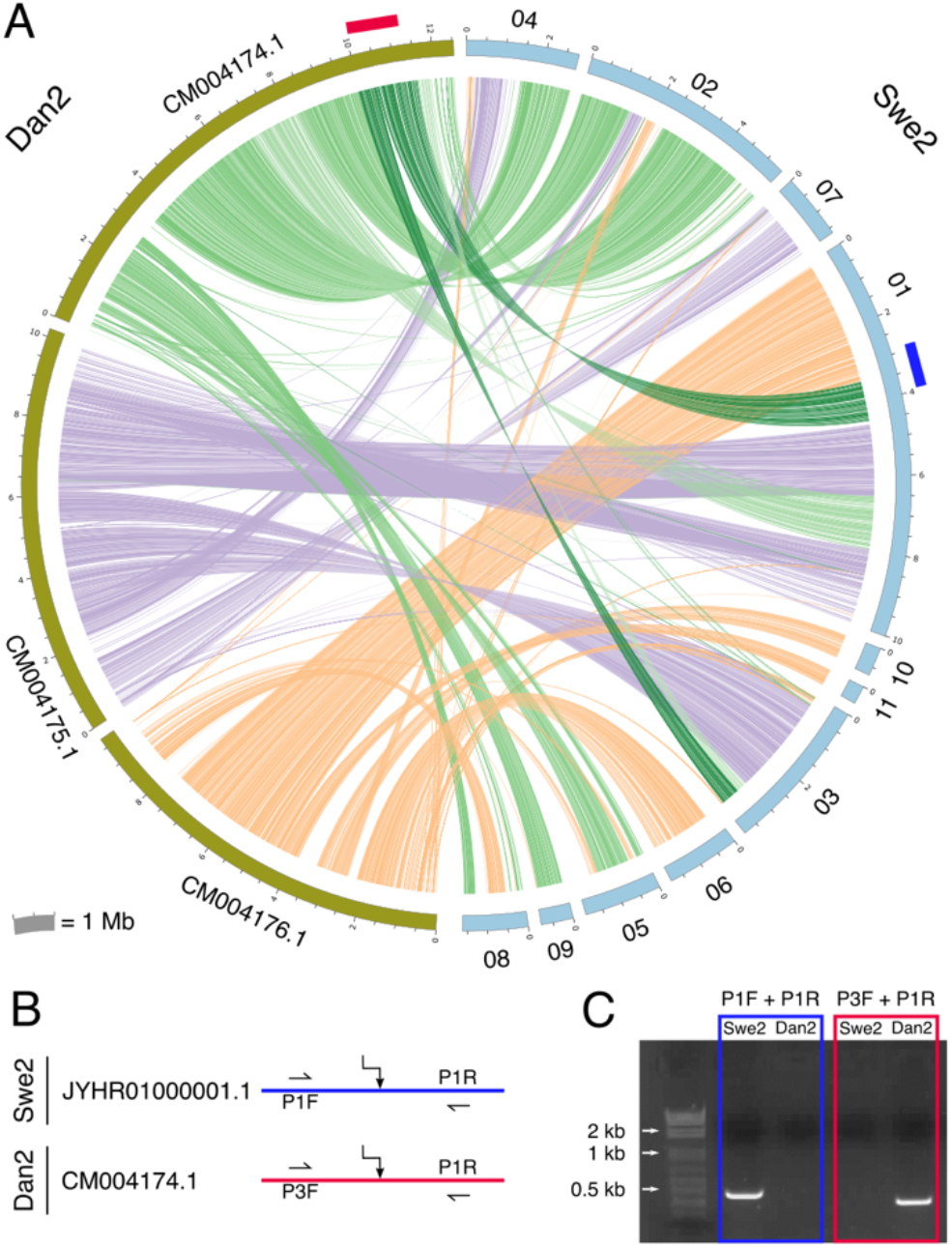
Inter-chromosomal rearrangements between strains Swe2 and Dan2. A. Circos plot representing regions >6 kb that are very similar between Dan2 (left – olive) and Swe2 (right – light blue) assemblies as determined by an all-against-all LAST analysis. Swe2 contig numbers are the last two digits of the accession ID (shown in B), preceding the suffix. Red and dark blue rectangles represent rearrangement junctions probed by PCR. B. Conceptual design of the PCR primers. C. Amplicons from the PCR were visualized after electrophoresis. Each pair gave one specific band of the expected size. The colour code is the same for the 3 panels.

All 3 nuclear genomes were divided in 3 similarly sized chromosomes, an unusual arrangement for such a fungus, as previously noted by Zhang *et al*. for Dan2 [6]. For the 2 strains related to Swe2, there was almost complete synteny of their nuclear genomes. Inspection of the one anomalous region in Swe1 where synteny broke down revealed that it was supported by only one long (215 kb) read and corresponded to a local discontinuity in the read coverage, as well as a break in the alignment between Canu-generated contigs and unitigs. All these factors indicated that this was an assembly artefact with a contig misassembled on the basis of an individual very long chimeric read (Supplementary Fig. S2). The same was true for the distinct unique non-syntenic region of the Swe3 assembly (Supplementary Fig. S3).

These were exceptional cases since the overwhelming majority of chimeric reads were identified and either trimmed or excluded from the assembly process by Canu (Supplementary Fig. S4, Supplementary Fig. S5). An in-depth analysis of the Swe1 chimeric reads revealed that a large proportion was in fact the consequence of sequencing errors. In almost 40% of cases (1010 / 2566), the two regions flanking the presumptive site of chimerism mapped to within 50 nucleotides of each other on the corresponding single scaffold. There was no discernible pattern to the distribution of this interval in the remaining candidate chimeric reads (Supplementary Fig. S6A-B), nor where there any regions that were more likely to be the site of chimeric junctions (Supplementary Fig. S6C).

Notably the single chimeric read that escaped censoring, leading to a misassembly of Swe1, was not identified by the dedicated tool YACRD, but was flagged as anomalous in reads recalled by Guppy (see Methods). This is an indication of the continuing improvement to base-calling tools. Also, these specific Swe1 and Swe3 misassemblies were absent from the corresponding chromosome assemblies produced by the *de novo* assembler Flye [8] (Figure 3A). This latter, however, introduced other assembly artefacts, including an erroneous fusion of contigs for the Dan1 assembly. This could not be ascribed to the inclusion of chimeric reads, but rather appeared to result from the incorrect treatment of repeat sequences, including telomeric repeats at the extremity of one of the fused contigs (Figure 3B-D). These results illustrate the interest of using more than one tool to aid in genome assembly. Therefore, starting with the Canu-generated sequences, we manually corrected anomalous regions and thereby produced assemblies for Swe1 and Swe3 that were entirely collinear (Figure 4A).

**Figure 3.**
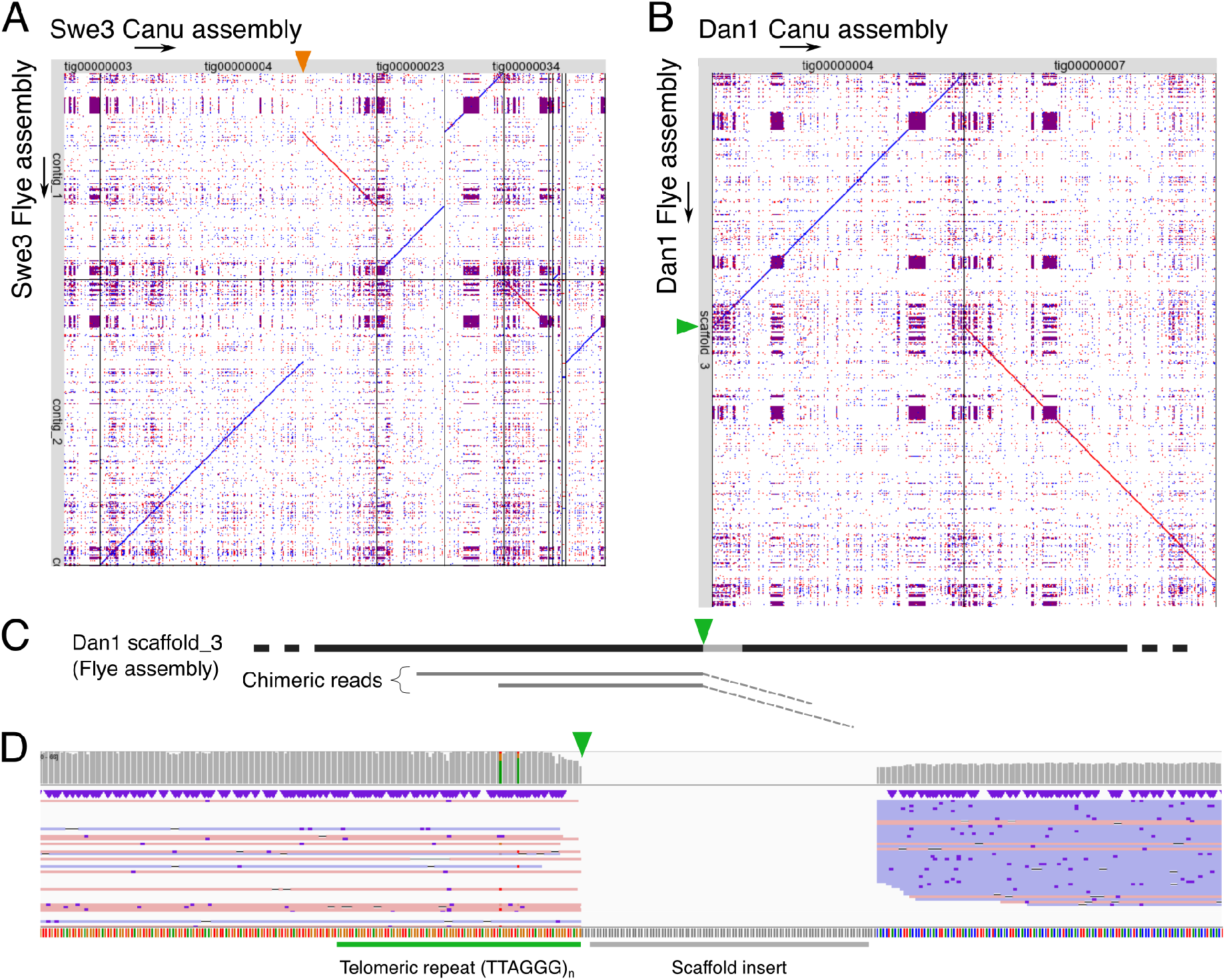
Comparisons between Canu and Flye assemblies. A, B. Dot plots of the non-congruent assemblies generated by Canu (x-axis) against those generated by Flye (y-axis) for the Swe3 (A) and Dan1 (B) genomes. The orange triangle (A) marks the position where the Canu contig *tig00000004* was split during the manual curation because of its chimeric nature. The green arrow (B) marks the position of a Flye scaffolding error. C. Schematic representation of the Dan1 Flye assembly, showing the mapping of chimeric reads close to the scaffolding error (green triangle). The coordinates in brackets are the mapping positions of the clipped part of the reads (dash line) on another contig of the assembly. Notably, this error was eliminated when these chimeric reads were excluded from the input data. D. Mapping of long-reads close to the scaffolding error (green triangle) on the Dan1 Flye assembly. The green bar marks the telomeric tandem repeat motif. The grey bar indicates the 100 Ns inserted by Flye to unite the scaffold.

**Figure 4.**
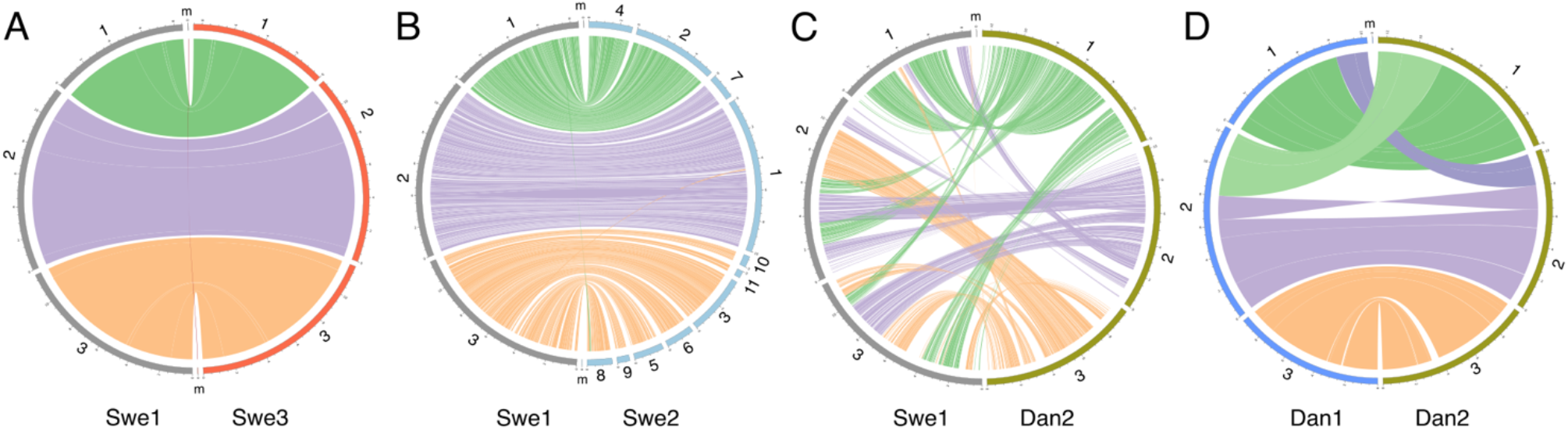
Synteny among the genomes of 5 *D*. *coniospora* strains. Circos plot representing regions >20 kb that are very similar between assemblies as determined by all-against-all LAST analyses. Each assembly is shown at the same scale and in the same order and orientation across panels.

These 2 genomes have 3 large chromosomes (8.5 Mb, 11.6 Mb, 11.6 Mb), each with identifiable telomeric [9] and centromeric regions, indicating that the overall genome structure has remained constant over 20 years of laboratory culture. This allowed us then to use the Swe1 sequence to scaffold the fragmented Swe2 genome (Figure 4B). To our great satisfaction, we were able to produce an entirely collinear chromosome scale assembly. Thus, it appears that there were no assembly errors in the published Swe2 genome, it was simply incompletely scaffolded. This applies equally to the genomic regions containing copies of some mitochondrial genes that we previous suggested might indicate assembly errors [5]. They were revealed to be accurate; *D. coniospora* has nuclear paralogous copies of 10 mitochondrial protein-coding and 15 tRNA genes (so called *numts* sequences [10]). These results give further support to the existence of long-term stability of the genome of the Swe2 related strains. A whole genome comparison between Swe1 and Dan2, however, revealed multiple and extensive genome rearrangements, involving intra- and inter-chromosomal translocations and inversions (Figure 4C).

Using the same strategy described above, we assembled and polished the Dan1 genome to give chromosome-level sequences. When we compared Dan1 and Dan2, we were surprised to find 2 major events of reciprocal exchange of chromosome ends, and an intra-chromosomal inversion (Figure 4D). These events were supported in a coherent and consistent manner by all the available data (Supplementary Fig. S7). In other fungal species, such chromosomal rearrangements have been reported to be the result of ectopic recombination between non-allelic homologous sequences, including repeated DNA elements [11,12]. A search of the 50 kb regions flanking each break point for transposable elements [13] and repetitive DNA families [14], failed to reveal any significant repeat sequence signature (see Supplementary Methods). As the Dan2 assembly is of high confidence, supported by long reads and optical mapping [6], given the short time of *in vitro* culture that separates it from Dan1, this suggests that the genome of the Dan1 isolate is not stable.

In alignments of the sequence of Swe1, generated using only nanopore reads, with that of Swe2, there were stretches of complete nucleotide identity extending over more than 25 kb. This is a testament to the general reliability of nanopore sequencing. We therefore identified the complete set of proteins identical in Swe2 and Dan2 corresponding to single copy, single exon genes (see Methods). These would be expected to be present in the newly assembled Swe1, Swe3 and Dan1 genomes. Indeed, using these 305 genes as a query, we could identify homologous sequences for each in all 3 genomes. Less than 1/6 of the corresponding genes, however, were predicted to encode full-length proteins in any of the 3 new genomes (Figure 5A). While nanopore reads are very useful for genome assembly, they suffer from a high error rate, especially in homopolymer stretches. Sequence quality can be improved using polishing tools that aim to ameliorate consensus sequences generally by going back to raw reads and applying integrative algorithms [15]. In our case, applying current best practices, while providing a very substantial improvement (up to 5-fold in the best case), did not take the prediction level beyond 82% accuracy. The quality of the prediction seen with the Dan1 genome was strikingly lower than the other 2 genomes (Figure 5A, Supplementary Table S1).

**Figure 5.**
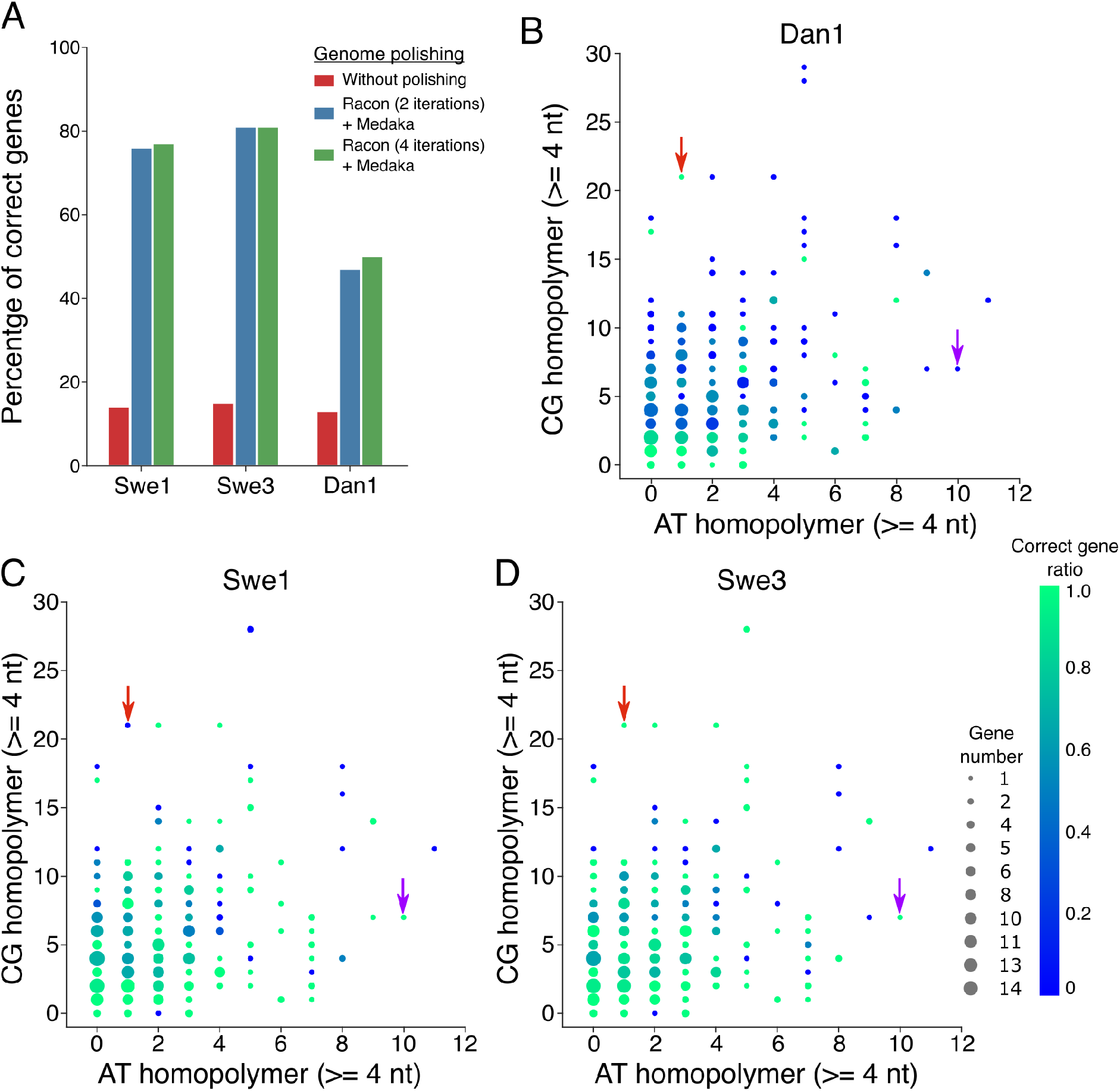
Evaluation of sequence errors in the 3 new genomes. A. Percentage of correct genes (based on length of the corresponding predicted protein) among 305 conserved genes, for the 3 new genomes, in the initial assembly and after two different polishing strategies. B, C, D. Scatter plots of homopolymer composition (A/T or C/G) and accuracy among the same 305 conserved genes for Dan1 (B), Swe1 (C) and Swe3 (D). The dot size is proportional to the number of genes, and the colour indicates the proportion of genes predicted to be correct. Red and purple arrows highlight two particular cases, among many, where homopolymer errors are only present in one genome.

Inspection suggested that the majority of errors were in homopolymer sequences, as expected, with nucleotide insertions and deletions leading to alterations of the reading frame. To investigate this poor homopolymer predictive performance systematically, we computed the number of G/C or A/T homopolymer stretches of at least 4 nucleotides for each of the 305 genes. We plotted these values, indicating the proportion of genes that encoded the expected full-length predicted protein for each of the 3 genomes. While there was the expected inverse relationship between accuracy and the number of homopolymer stretches, there were striking exceptions. Curiously some of these exceptions were specific to a single genome (Figure 5B-D). Further, and unexpectedly, polishing introduced more nucleotide insertion errors than deletions, frequently on the basis of tenuous read support. Overall, however, there was no obvious pattern to explain why errors were introduced, given the underlying reads used to build the consensus sequence (Supplementary Table S2).

During the inspection of the assembled and polished genomes, we found two other types of anomalies. The first concerned the regions flanking the nuclear genomic copies of mitochondrial genes (*numts*), where polishing added short extraneous low complexity sequences (average length 15 nt, mainly As or Ts), for which, surprisingly there was no sequence support from the reads used by the assembler (Figure 6A). This probably arose because of the very high nucleotide similarity between regions of the nuclear and mitochondrial genomes that extended across more than 25 kb, including a repeat of 9.8 kb (Supplementary Fig. S8A-B). Notably, despite using high coverage ONT long reads, we could not establish with absolute certainty the precise copy number for the unit sequence in the Swe genomes (Supplementary Fig. S8B).

**Figure 6.**
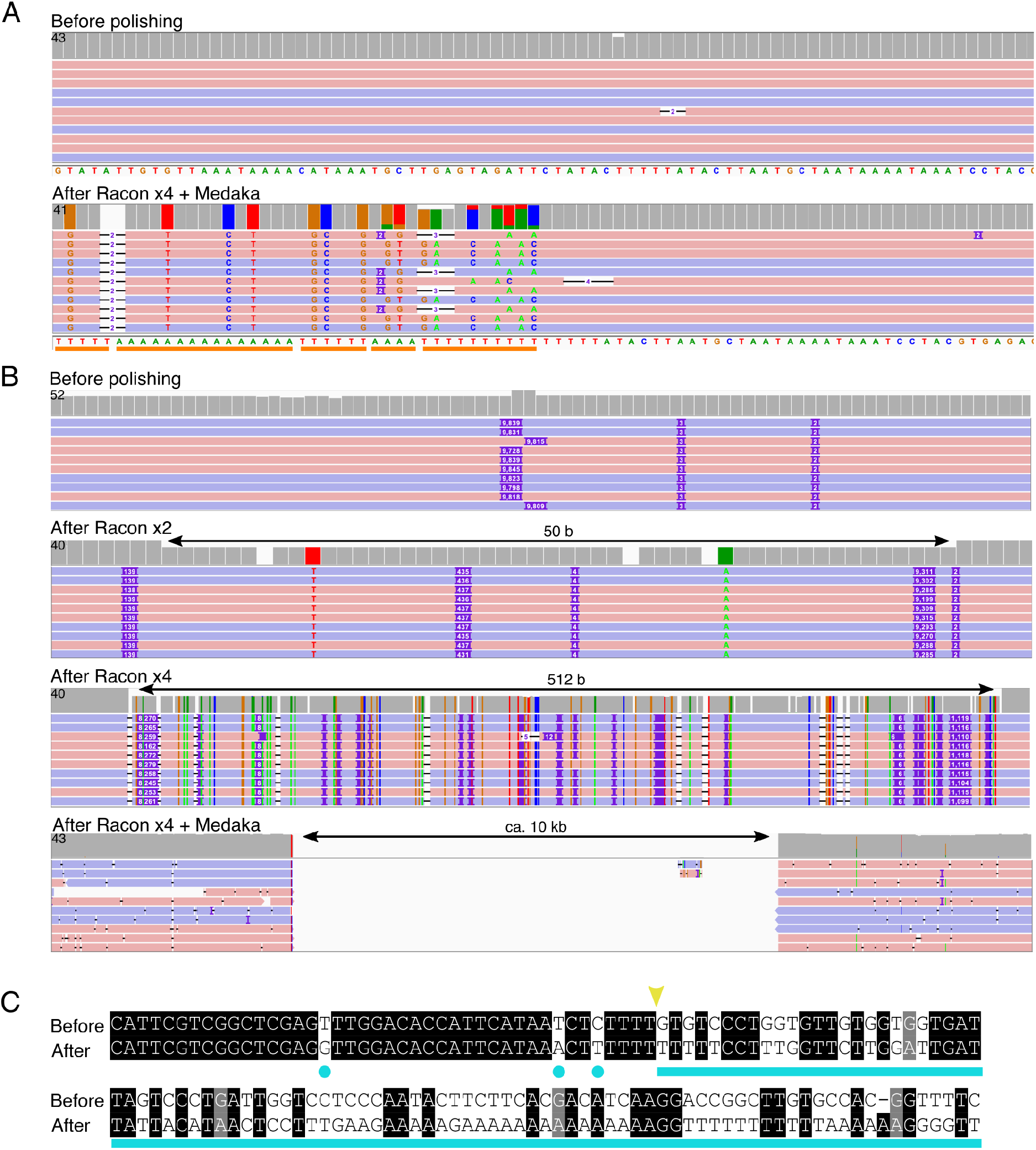
Sequence anomalies introduced by assembly and/or polishing tools. A. A comparison of one small region of the Swe3 sequence before (top) and after polishing (Racon x4 and Medaka; bottom). As indicated by the orange line, long stretches of A and T homopolymers are introduced by polishing, in the absence of coherent read support. B. From top to bottom, the assembly produced by Canu excludes a region of around 10 kb, despite strong read support. After 2 and 4 iterations, Racon progressively filled the gap. Medaka then introduced an insert of roughly the correct size, but of aberrant sequence composition. For each panel, the height of the boxes in the top line indicates the read coverage for each base. A grey box indicates full agreement with the consensus sequence, otherwise the colour indicates the proportion of read support for each nucleotide (G, tan; C, blue; A, green; T, red). Below this, the ONT reads that align in forward (pink) and reverse (blue) orientation are shown as lines. A coloured letter or purple rectangle show a difference (nucleotide variant or insertion in reads, respectively) in the read’s sequence compared to the genome sequence. C. The 10 kb sequence introduced by polishing is of aberrant composition as illustrated by the region immediately surrounding the 5’ breakpoint (yellow arrowhead). There are single nucleotide errors introduced despite coherent read support for the “Before” sequence (light blue dots), and then a continuous stretch, exemplified by A and T homopolymers that lack any sequence support at all (light blue line).

In the second case, for the Swe3 genome, a large (ca. 10 kb) region, with a complex sequence, well supported by the Canu corrected and trimmed reads, was inexplicably excluded from the initial Canu assembly and only imprecisely restored by polishing (Figure 6B-C, Supplementary Fig S8C). Here, while there was no evidence for repeated DNA elements on both sides of the point of sequence discontinuity, there was a single such 1.2 kb duplication (Supplementary Fig S8C-D). These few regions were identified because of discontinuities in the depth of read coverage, which otherwise was remarkably constant across the complete genomes. With the resolution of these assembly errors, we were able to generate complete genomes of high overall structural quality using ONT long reads only.

As explained above, however, these assemblies were not of sufficient sequence quality to allow accurate gene prediction. Therefore, to extend our analysis, we used Illumina sequencing to generate very deep short read coverage for the Swe1, Swe3 and Dan1 genomes. This allowed high quality final sequences to be generated for all 3 strains. While short-read-based polishing did not alter the global structure, it allowed homopolymer length errors to be corrected and the generation of entirely contiguous chromosome sequences (Supplementary Table S1).

To confirm the correctness of the short-read polished assemblies, we returned to our 305 single copy orthologues. After the short-read polishing, all 305 genes could be identified in each of the 3 genomes (Supplementary Table S1). We also benchmarked our successive assemblies using BUSCO that searches for a set of universal single-copy orthologues (USCO) by sequence similarity. While the initial genome assemblies gave low scores, with roughly 65% of complete USCOs and 35% fragmented or missing (Table 1), after long-read polishing the score for complete USCOs increased up to as high as 97%. Given the demonstrably low quality of the genome sequences (Figure 5), we investigated the basis of this disparity. We identified among the USCOs those that corresponded to single exon genes in the Dan2 and Swe2 reference genomes. These genes were then used as queries for high-stringency searches of the Dan1, Swe1 and Swe3 genomes at successive steps of assembly and polishing and the results compared to the results of the corresponding BUSCO analysis. While BUSCO gave no false negatives, it gave a large number of false positives, except in the analysis of the short-read polished genomes (Figure 7A). These arose because BUSCO was not sufficiently sensitive to the presence of short indels. As an example, the Swe1 gene corresponding to RJ55_06485 had the expected sequence after short-read polishing. Two errors in homopolymer sequences led to 2 frameshifts in the unpolished assembly. One of these was corrected by long-read polishing, but for the other there was an over-compensation, leading to a different frameshift (Figure 7B). In both assemblies, these errors were compatible with open-reading frames that collectively reconstituted a close ortholog of RJ55_06485 leading to the erroneous BUSCO result. As discussed below, this analysis highlights the fact that BUSCO scores based on sequence alignments are not an appropriate measure for ONT-only eukaryotic genomes. The BUSCO score rose to nearly 99% after the short-read polishing. In this case, the figures accurately reflect genome completeness and quality (Figure 7A). These figures are comparable to those for the previous Dan2 and Swe2 assemblies. The new Swe1, Swe3 and Dan1 genomes therefore represent the starting point for future detailed analysis to characterise the molecular evolution of *D. coniospora*.

**Table 1:** BUSCO results

| Strain | Assembly | Complete | Complete: single | Complete: duplicated | Fragmented | Missing |
| --- | --- | --- | --- | --- | --- | --- |
| Dan1 | Canu curated | 820 (62.4%) | 820 (62.4%) | 0 (0%) | 259 (19.7%) | 236 (17.9%) |
| Dan1 | Long-read polish | 1187 (90.3%) | 1187 (90.3%) | 0 (0%) | 62 (4.7%) | 66 (5%) |
| Dan1 | Short-read polish | 1297 (98.6%) | 1296 (98.6%) | 1 (0.1%) | 9 (0.7%) | 9 (0.7%) |
| Dan2 | [6] | 1298 (98.7%) | 1297 (98.6%) | 1 (0.1%) | 8 (0.6%) | 9 (0.7%) |
| Swe1 | Canu curated | 869 (66.1%) | 868 (66%) | 1 (0.1%) | 243 (18.5%) | 203 (15.4%) |
| Swe1 | Long-read polish | 1266 (96.6%) | 1266 (96.6%) | 0 (0%) | 21 (1.6%) | 28 (2.1%) |
| Swe1 | Short-read polish | 1296 (98.6%) | 1295 (98.5%) | 1 (0.1%) | 8 (0.6%) | 11 (0.8%) |
| Swe2 | [5] | 1296 (98.6%) | 1294 (98.4%) | 2 (0.2%) | 9 (0.7%) | 10 (0.8%) |
| Swe3 | Canu curated | 859 (65.3%) | 858 (65.2%) | 1 (0.1%) | 243 (18.5%) | 213 (16.2%) |
| Swe3 | Long-read polish | 1274 (96.9%) | 1274 (96.9%) | 0 (0%) | 17 (1.3%) | 24 (1.8%) |
| Swe3 | Short-read polish | 1295 (98.5%) | 1294 (98.4%) | 1 (0.1%) | 9 (0.7%) | 11 (0.8%) |

Percentage of each category of the expected 1315 USCOs for different genome assemblies. Of the 11 USCOs missing in Swe1 and Swe3, 10 are also absent from Swe2, and 9 from Dan1 (and Dan2). These are therefore likely to be real gene losses in *D. coniospora*, so that only 2 USCOs (0.2%) at most are missing.

**Figure 7.**
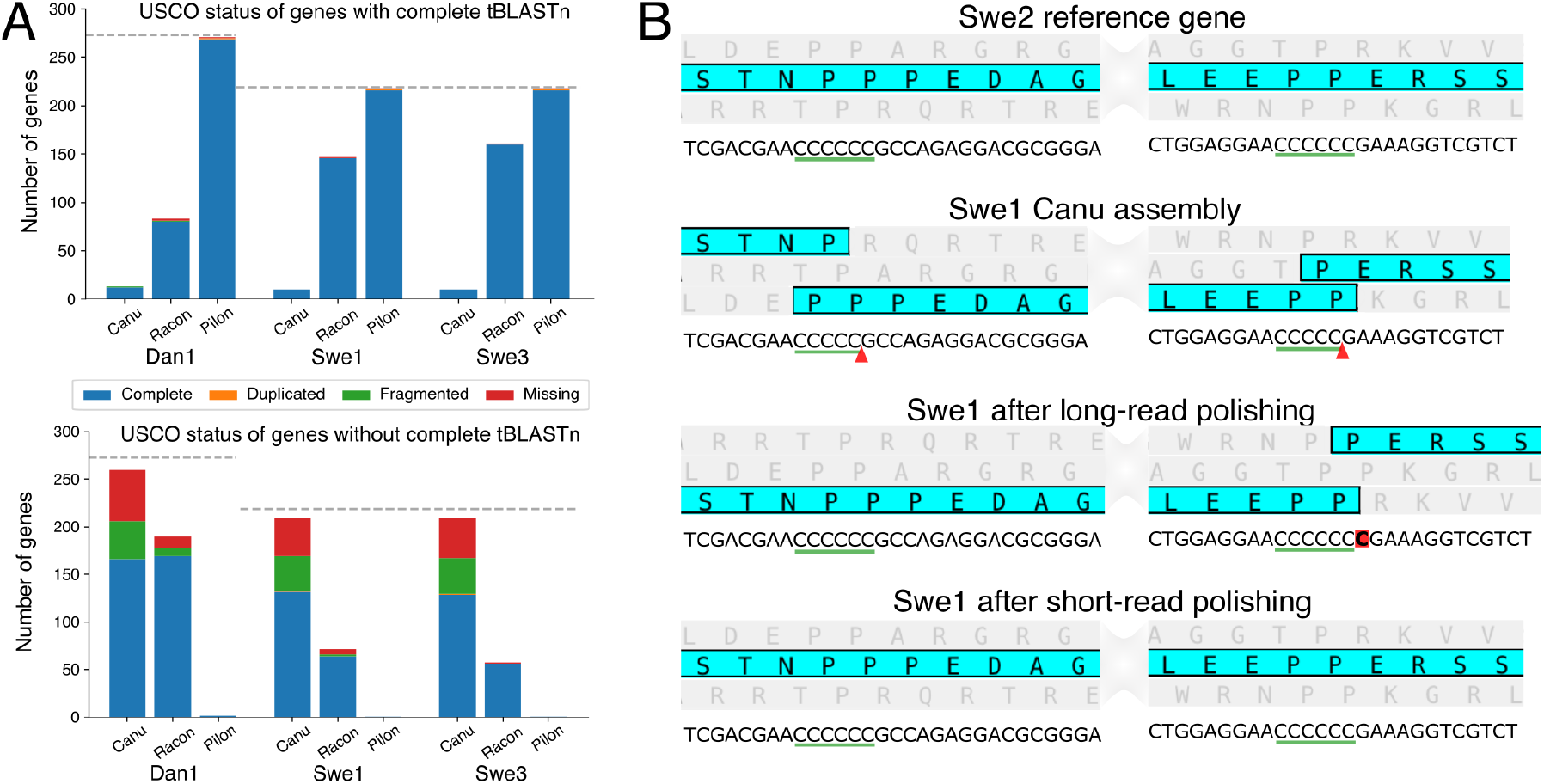
Example of sequence errors introduced during assembly and polishing. A. Stacked bar plot of USCO status for the orthologues of selected mono-exonic Swe2 or Dan2 genes classified according to the result of a TBLASTN search against the indicated assembly (Canu: from Canu; Racon: after long-read polish; Pilon: after short-read polishing). B. Detailed view of 2 parts of RJ55_06485 from the Swe2 reference genome each containing a homopolymer sequence (underlined) and the corresponding positions in successive Swe1 assemblies. For each, the predicted protein sequence, highlighted in turquoise, with the other open reading frames in grey, is shown above the corresponding nucleotide sequence. The red arrow heads highlight the missing nucleotides, the extraneous nucleotide is boxed in red.

## Discussion and conclusion

Previous genome assemblies for *D. coniospora* required a combination of sequencing approaches [5,6]. Here, using only long reads and Canu, we produced the first complete circular mitochondrial genome for *D. coniospora* and were able to generate chromosome-scale assemblies for the nuclear genome. The rare misassembled contigs, formed by Canu because of single very long chimeric reads, as previously described [16], could be detected by read coverage anomalies and comparisons with unitigs, suggesting that solutions to avoid their creation could be implemented within Canu. The majority of reads that were flagged as chimeric arose from sequencing or polishing errors. They reflected a short (<50 bp) discrepancy between the individual reads and the final sequence. There was no indication of any sequence bias at the break points of the remaining chimeric reads supporting the notion that these reads arise from too rapid reloading of the sequencing pore [17].

The use of other genome assembly tools, and the comparison of assembly discrepancies is an additional method to produce high confidence genomes. Here, we used Flye that for these genomes required run times that were ten-fold shorter than Canu. A comparison of the assemblies highlighted ambiguous regions in the genome that could then be resolved by manual inspection. On the other hand, Flye was confounded by telomeric repeats. Since telomeres can be identified on the basis of their sequence, there is also clear room for algorithmic improvement to Flye through the explicit definition of chromosome ends.

One clear and well-established advantage of using long reads is the possibility of resolving very extended stretches of complex tandem repeats (VeCTRs) [18] and other repetitive sequences including centromeres. These correspond to most of the breaks in the continuity of the published Swe2 genome. In addition to acrocentric regional centromeres, Zhang *et al*. reported the presence of a vestigial centromere from a putative chromosomal fusion event [6]. These were also found in the fully assembled Swe1 and Swe3 genomes, indicating that chromosomal fusions were present in the common ancestor of the Swe1 and Dan1 strains.

For Swe1, Swe3 and Dan1 we were able to reconstruct complete mitochondrial genomes, with features typical of fungi of the order Hypocreales. On the other hand, unlike Dan1 (and Dan2), the nuclear genomes of Swe1 and its derivatives Swe2 and Swe3, contained different numbers of copies of sequence very similar to parts of their own mtDNA. This type of event, and more generally repeated regions with long and nearly identical sequences are more readily detectable with long reads [19], and are particularly challenging for polishing even with short reads [20].

The duplication of mitochondrial genes in the nuclear genome has been described in other fungal genomes [10] and must have occurred after the divergence of Dan1 and Swe1. Despite this genome plasticity, even after 20 years of continuous laboratory culture the Swe1 and Swe3 genomes were entirely collinear. This contrasts with the rearrangements seen between the Dan1 and Dan2 genomes that in principal should be from strains that have had little opportunity to diverge (L. Castrillo, Curator, ARS Collection of Entomopathogenic Fungal Cultures, personal communication). It will be interesting in the future to characterize the reasons for the marked difference in genomic stability between Dan1 and Swe1.

The accuracy of ONT long read sequencing is increasing because of improvements in the chemistry used, signal detection, as well as base-calling [21]. Despite good read depth, however, our assemblies were not of sufficient quality at the nucleotide level to allow accurate gene prediction. Further, we noted that although polishing using only long reads dramatically increased overall sequence accuracy, it introduced errors around the *numts*. Similar errors during polishing of near identical sequences has been noted in ONT-based metagenomic studies [22]. Despite these limitations, research groups are publishing and submitting to public sequence databases genomes for fungi, plants and animals based on nanopore sequencing alone (86 for Eukaryotes in addition to the 134 Bacterial genomes in “Assembly” from GenBank release 236 from the 2020/02/15). This is problematic as low-quality genome sequences compromise the accuracy of sequence similarity searches in public databases. On the basis of our results, a re-analysis of the completeness of these “nanopore-only” genomes is merited, to confirm that they are indeed low quality. Similar concerns do not apply to fungal genomes assembled using only long reads generated with Pacific Bioscience technology [23] as these do not suffer from the intrinsic problem of homopolymer length errors that we found to be the most significant quality barrier when using ONT reads. On the basis of our detailed analysis and in line with the consensus regarding *de novo* assembly with ONT long reads (e.g. [24]), we polished our 3 assemblies with short reads. This greatly improved their quality.

Regarding the homopolymer sequence errors, as noted above, they were not consistent across the sequenced genomes; even between Swe1 and Swe3 there were instances of widely differing rates of errors in orthologous genes, despite very similar underlying reads. Indeed there was no clear pattern in the inaccuracies, which will render bioinformatics approaches to remedy this problem more difficult. On the other hand, the errors were more often over-prediction of homopolymer length, despite having a majority of reads supporting the correct sequence. It is possible that polishing tools have not kept pace with improvements in base-calling, leading to an over-compensation in the inference of homopolymer length.

It is standard practice to check the completeness of *de novo* genome assemblies with a strategy based on the detection of predicted groups of conserved orthologous proteins. One popular and much cited tool is BUSCO [25] which was developed before ONT-based sequencing became prevalent. Since BUSCO relies on *in silico* translation, small indels can be overlooked as the resulting virtual sequence can be recapitulated despite a frameshift. This explains the disparity between the BUSCO results and our own analyses that were deliberately restricted to mono-exonic genes. Contrary to BUSCO, our analysis indicated that about 1/5 of the genes after long-read polishing had an incorrect sequence. Current BUSCO-type approaches, based on sequence similarity and not excluding genes with improbably short introns, cannot be used as a quality metric for ONT-only assemblies, and are appropriate only after short-read correction.

In conclusion, nanopore long read sequencing provides a powerful way to assemble complex genomes with limited manual curation but still fall short of the quality required to produce publishable eukaryotic genomes. In our case, it has revealed new information about genome plasticity in *D. coniospora* and provided a backbone that will permit future detailed study to characterize gene evolution in this important model fungal pathogen.

## Methods

### DNA extraction

*D*. *coniospora* spores were cultured in liquid NGMY medium [26] at 37°C for 5 days. Fungal DNA was extracted according to a published protocol (from p13 onwards of [27]) [28], with the following modifications: instead of centrifugation to collect DNA after precipitation with isopropanol, we recovered the DNA filaments with a glass hook, washed and dried them as described [29] and resuspended the DNA without agitation in Tris-EDTA buffer.

### Nanopore sequencing library preparation

Libraries were prepared for sequencing on GridION with the ligation sequencing kit SQK-LSK109. The GridION sequencing was run on flowcell FLO-MIN106 for 47, 48 and 48 hours using 972, 660 and 610 ng of DNA (for Swe3, Swe1, Dan1 respectively) and MinKNOW 2.1 v18.06.2.

### Illumina sequencing library preparations

The same DNA samples were used to prepare paired-end libraries with insert size of circa 680 bp, following the manufacturer’s instructions for the kit NEBNext® Ultra™ II DNA (New England Biolabs Inc. Ipswich, MA, USA). The libraries were sequenced using an Illumina NextSeq500 system (s/n NB501764).

### Basecalling, adaptor trimming and chimeric read detection

For a first assembly, reads were basecalled at the EMBL using Guppy v1.5.1 (Oxford Nanopore Technologies). For subsequent polishing, we used Guppy v3.0.3 (with parameters -c dna_r9.4.1_450bps_hac.cfg), then adaptors were trimmed with Porechop v0.2.4 [30] with default parameters. YACRD v0.5.1 [31] with the subcommand chimeric and the option --filter was used to remove chimeric reads.

### Whole genome alignments

Genomes were aligned using LAST v979 [32]. A database was first generated (last-db -cR01), and then lastal and last-dotplot with default parameters were used to generate respectively an alignment file and a dot-plot. For the circular visualization of genome alignments, we used the command lastal with -f BlastTab parameter, then parsed the alignment to filter out short alignments and generate the links file needed by Circos [33].

### Mapping of long reads

Validation of genomes during and after assembly involved rounds of read mapping. Reads were aligned with Minimap2 v2.16r922 [34] (with parameters -ax map-ont). The resulting mapping file was processed with Samtools v1.9 [35] to obtain a sorted BAM file (samtools view -bS -q 1 -F 4; samtools sort; samtools index). Mapping results were visualized with IGV v2.5.0 [36].

### Genome assembly

Assemblies were performed with Canu v1.7 [37] and the parameters useGrid=False, genomeSize=30m, correctedErrorRate=0.16 with reads basecalled by Guppy v1.5.1. For the manual curation of the assemblies, we generated whole assembly alignments and dot-plots of Swe1, Swe2 and Swe3 two by two. For Swe1 and Swe3, Canu contigs were ordered by synthesizing the results from the 3 possible all-against-all alignments. To confirm a link between two contigs, we employed the following strategy: when a contig of the Swe1 assembly spanned two contigs of Swe3, long reads of Swe1 present in this spanning area were extracted from the Swe1 corrected and trimmed reads provided by Canu. Then this set of reads was mapped on Swe2 and Swe3 assemblies. The two targeted contigs of Swe3 were considered ‘linked’ if different parts of several unique reads mapped on the two Swe3 contigs ends. If the reads that supported the link had different mapping orientation (forward or reverse), one contig was complemented before the last step (see *Solving links between contigs*) to ensure a correct orientation of the final chromosome.

To guide correct assembly, we also searched for centromeres in the contigs. They were identified as highly duplicated regions in the all-against-all alignment dot-plots produced by LAST. The identification of the repeated canonical telomeric sequence (TTAGGG)_n_ [9] and its reverse complement (CCCTAA)_n_ at the beginning or end of certain contigs allowed the identification of chromosome ends. The Dan1 assembly was manually curated using a similar strategy with the Dan2 genome as a reference.

### Solving links between contigs

Overlaps between linked contigs were identified by a BLASTN [38] alignment of their last 100 kb. Any duplicate sequence was trimmed out from one contig and both contigs were joined. The inferred junction was then validated by verification of the underlying read support. For the linked contigs that did not overlap, the sequence in the gap was extrapolated from the reads that matched and extended the ends of contigs, on the basis of alignments at the last 1 kb of each contig. These sequences were aligned with MAFFT v7.427 [39]. The alignment was visualized with SeaView [40], and only the portion of the alignment strictly between the two contigs sequences was kept. Seaview also generated a consensus sequence (on the basis of 60 % sequence identify by default). The resulting sequence was inserted between the two contigs to link them and the supposed continuity verified by a further cycle of read mapping.

### Assembly polishing with long-reads

Genome polishing was carried out with 2 or 4 iterative runs of Racon v1.4.2 [41] and parameters -m 8 -x -6 -g -8 -w 500, and a run of Medaka v0.8.1 (Oxford Nanopore Technologies) with the parameter -m r941_min_high.

### Mitochondrial genome circularization

Canu assembles small circular elements as contigs with tandem duplications of the element. We resolved the mitochondrial genomes as recommended by Canu’s authors [7]. MUMmer suite v4.0.0.beta2 [42] was used to align the contig identified as the putative mitochondria on itself with NUCmer and parameters --maxmatch --nosimplify. Coordinates of a full copy were identified with the show-coords command and -lrcT parameters.

### PCR

PCR was carried out to test a genome rearrangement between Swe2 and Dan2 genomes, with primers P1F (GAGATATCGAACGTCGCATGG), P1R (ACATCAAGCCTTTGTCGAGGA), and P3F (GCTCAGGACCGACGTACAAG). PCR reactions were run according to the GoTaq® G2 Flexi DNA polymerase instructions (Promega), with 50 ng of template DNA, 1 mM of each forward and reverse primer, in a final volume of 25 *μ*L. The reaction started by initial denaturation at 95°C for 2 min, followed by 30 amplification cycles (95°C for 30 sec, 60°C for 30 sec and 72°C for 30 sec), and a final elongation for 5 min at 72°C.

### Defining a set of 305 identical proteins

Identical proteins shared by the two *D.* coniospora genomes available (Swe2 and Dan2) were recovered using a reciprocal best BLAST [38] hit strategy on the two proteomes. Proteins that were duplicated in one or both genomes were filtered out. The set was further refined by only retaining proteins corresponding to mono-exonic genes.

### Assessment of gene sequence in ONT-only assemblies

TBLASTN searches were run using the amino-acid sequence of the set of 305 identical proteins against the different nanopore only assemblies. A gene was considered as correct if the query coverage, *i.e* the ratio of alignment length over the query length, was equal to 1.

### Short read polishing

Shorts reads were trimmed using Trimmomatic v0.39 [43] with the parameters LEADING:3 TRAILING:3 SLIDINGWINDOW:4:30 MINLEN:36. Then, only paired reads were mapped on assemblies with bwa v0.7.17 [44] and default parameters (bwa index, then bwa mem). The resulting mapping file was converted in BAM, sorted and indexed with samtools. This latter file was used to polish the assembly with Pilon v1.23 [45] with the parameters --fix bases --vcf --mindepth 10 --minmq 20 --minqual 15 --changes --diploid. Several iterations were conducted for each strain, until the number of changes was less than 5.

### Flye assembly

An additional *de novo* assembly was performed with Flye v2.4.2 [8], and the parameter --genome-size 32m, using the ONT reads recalled by Guppy v3.0.3.

### Assessing the genome integrity

The genome integrity was assessed with BUSCO v3.1.0 and the curated set *ascomycota_odb9* version 2016-02-13 [25]. A BLASTP search enabled Swe2 monoexonic genes present among USCOs to be identified. This list of 219 Swe2 genes was then used as a TBLASTN query against the different assemblies of Swe1 and Swe3. A gene was considered correct when it matched the corresponding Swe2 gene perfectly in length. An analogous analysis was carried out for Dan1, on the basis of the 273 Dan2 monoexonic genes that are USCOs.

### Characterisation of chimeric reads

Swe1 reads identified as chimeric by YACRD were aligned on the final (short-read polished) Swe1 assembly. The main alignment was identified using samtools view -F 2308. The CIGAR string was then parsed to determine whether the longest residual part of the read was 5’ or 3’ to the main alignment, thereby giving an orientation to the putative chimeric read and localising the potential chimeric break point. The 500 bp of sequence 5’ and 3’ of this point were extracted and individually mapped back on the Swe1 final assembly and the number of unique reads in a 10 kb non-overlapping sliding window was calculated. For the reads for which both 500 bp fragments mapped on the same chromosome, the smallest distance between the two fragments was calculated.

## Data availability

Genomes of the strains Swe1, Swe3 and Dan1 are available on our institute website (http://www.ciml.univ-mrs.fr/applications/DC/Genome.htm). All the reads used in this work can be found at the European Nucleotide Archive (ENA) under the study numbers PRJEB35969, PRJEB35970 and PRJEB35971. The raw signal runs are available under the accessions ERR3774158, ERR3774162 and ERR3774163; the FASTQ files of basecalled reads (Guppy v3.0.3) are available under the accessions ERR3997391, ERR3997394 and ERR3997483; the FASTQ files of Illumina paired-end reads are available under the accessions ERR3997389, ERR3997392, ERR3997395. Accession numbers are given in the order Swe1, Swe3 and Dan1.

## Acknowledgments

The authors thank Yuquan Xu and Liwen Zhang for providing access to the raw sequencing data for Dan2, Xing Zhang for help in preparing DNA samples, Lionel Spinelli for informatic support and Nathalie Pujol and Laurent Tichit for comments.

## Supplementary data

*supplementary_figures.pdf* contains 8 supplementary figures.

*supplementary_table_1.xls contains* the read coverage of the genomes. This table also contains the results for the TBLASTN on the 305 candidate identical proteins.

*supplementary_table_2.xls* is a table recording read support, in Swe1, Swe3 and Dan1 assemblies, for the predicted correct sequence for each homopolymer stretch in the genes corresponding to 10 protein of the 305 candidate identical proteins.

*supplementary_methods.pdf* contains additional methodological details.

---

**Supplementary Figure 1.**
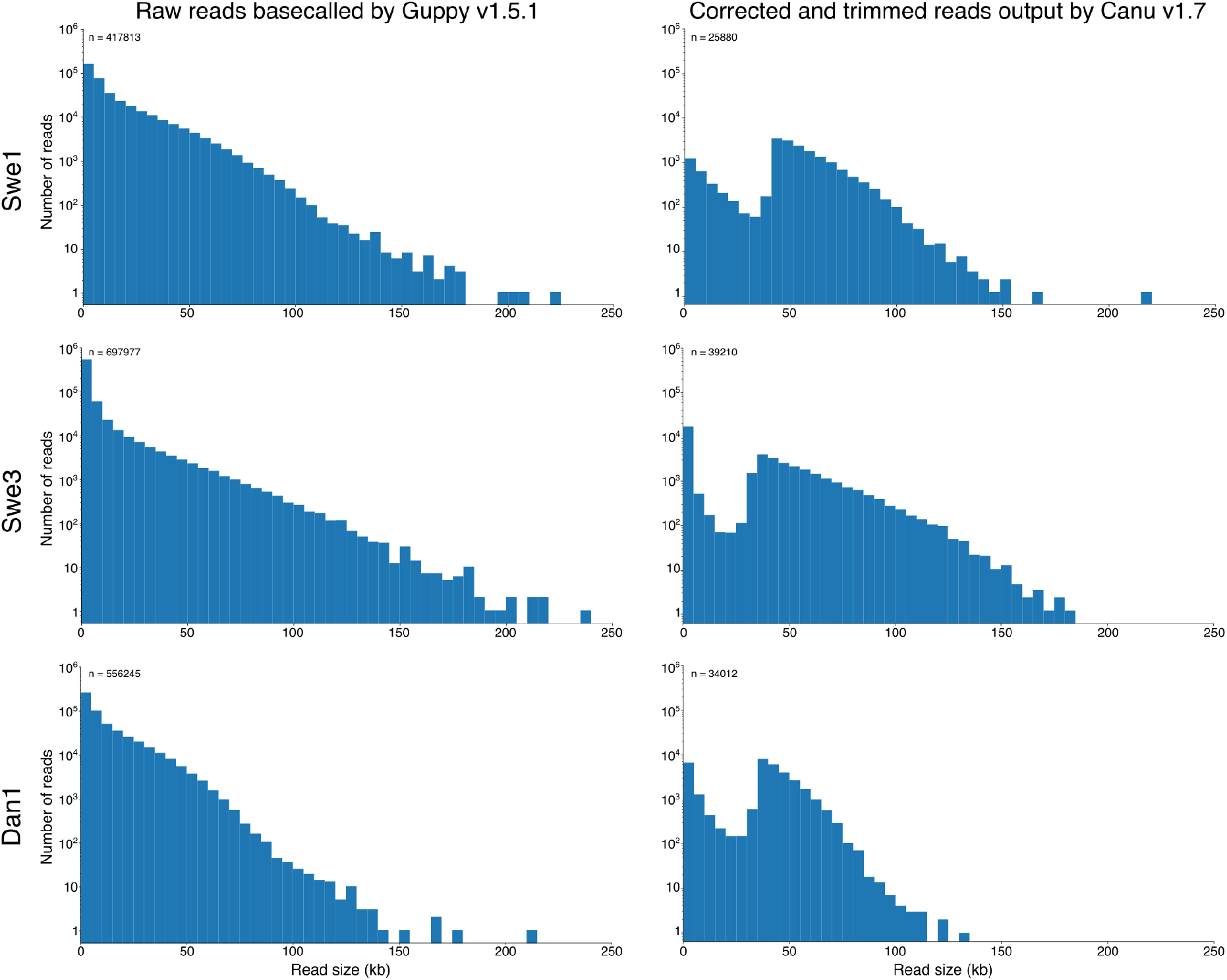
Distribution of the size of the reads and the assembly statistics. Distributions in 5 kb bins of the size of the set of reads basecalled by Guppy v1.5.1(left-hand panels), and of reads corrected and trimmed by Canu for the initial assemblies (righthand panels).

**Supplementary Figure 2.**
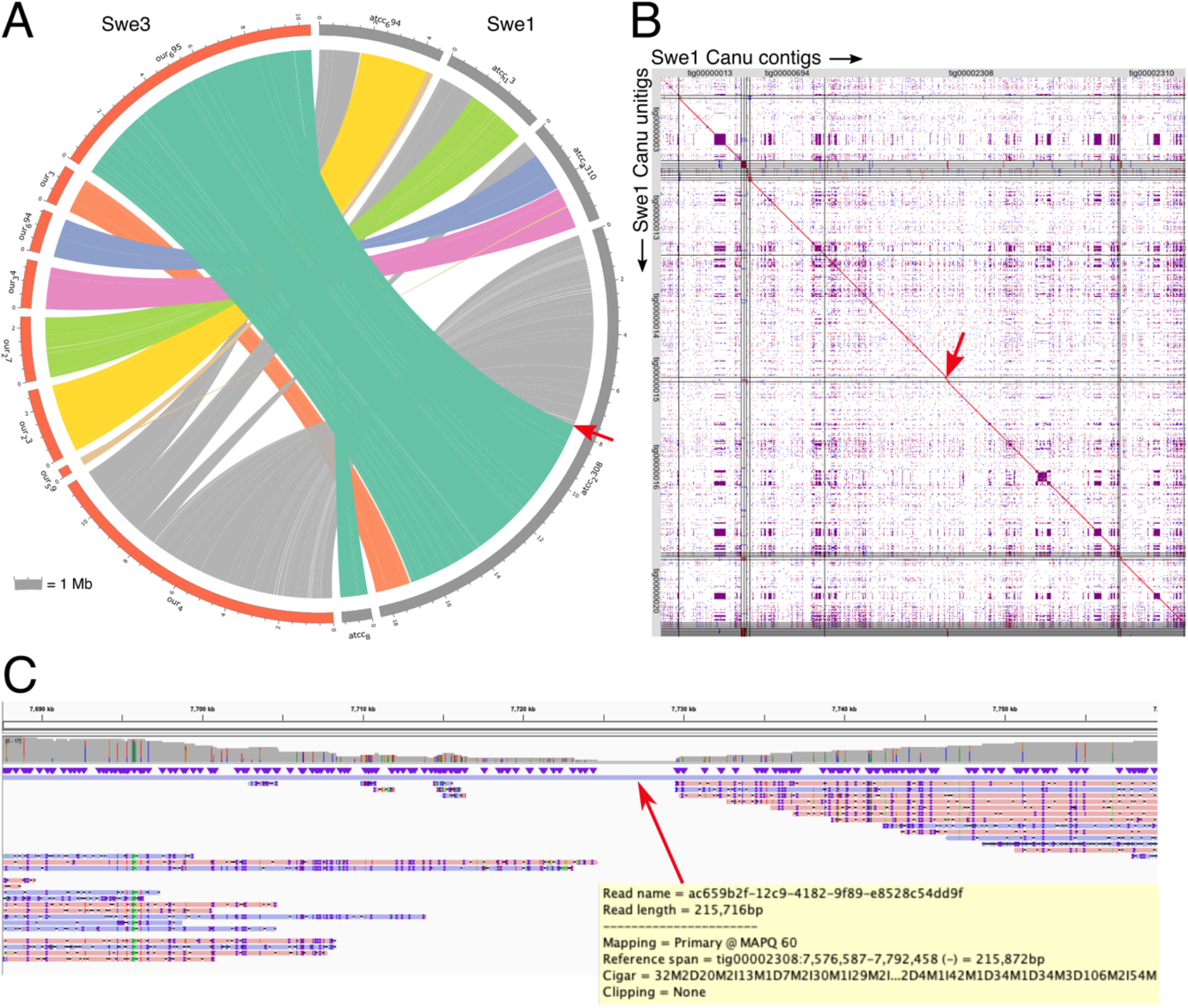
Detection of an error in the initial Swe1 assembly introduced by a long chimeric read. A. Circos plot representing regions >20 kb that are very similar between Swe3 and Swe1 Canu assemblies as determined by an all-against-all LAST analysis. The red arrow indicates a break in the synteny for the largest Swe1 contig. B. Dot-plot of an all-against-all comparison of the Swe1 contigs and unitigs produced by Canu. Contigs are contiguous sequences present in the primary assembly, including both unique and repetitive elements. Unitigs are contigs split at alternate paths in the assembly graph. The red arrow indicates the discontinuity in the alignment between Swe1 contigs and unitigs. This occurs on the same contigs and at the same coordinates as in A. C. Mapping of the Swe1 reads, corrected and trimmed by Canu, on the Swe1 Canu assembly (detail of around 70 kb on contig tig00002308 flanking the synteny break). Moving into the central 5 kb region, read support progressively drops from 40 to just one, corresponding to a very long read of about 215 kb, which was shown to be chimeric.

**Supplementary Figure 3.**
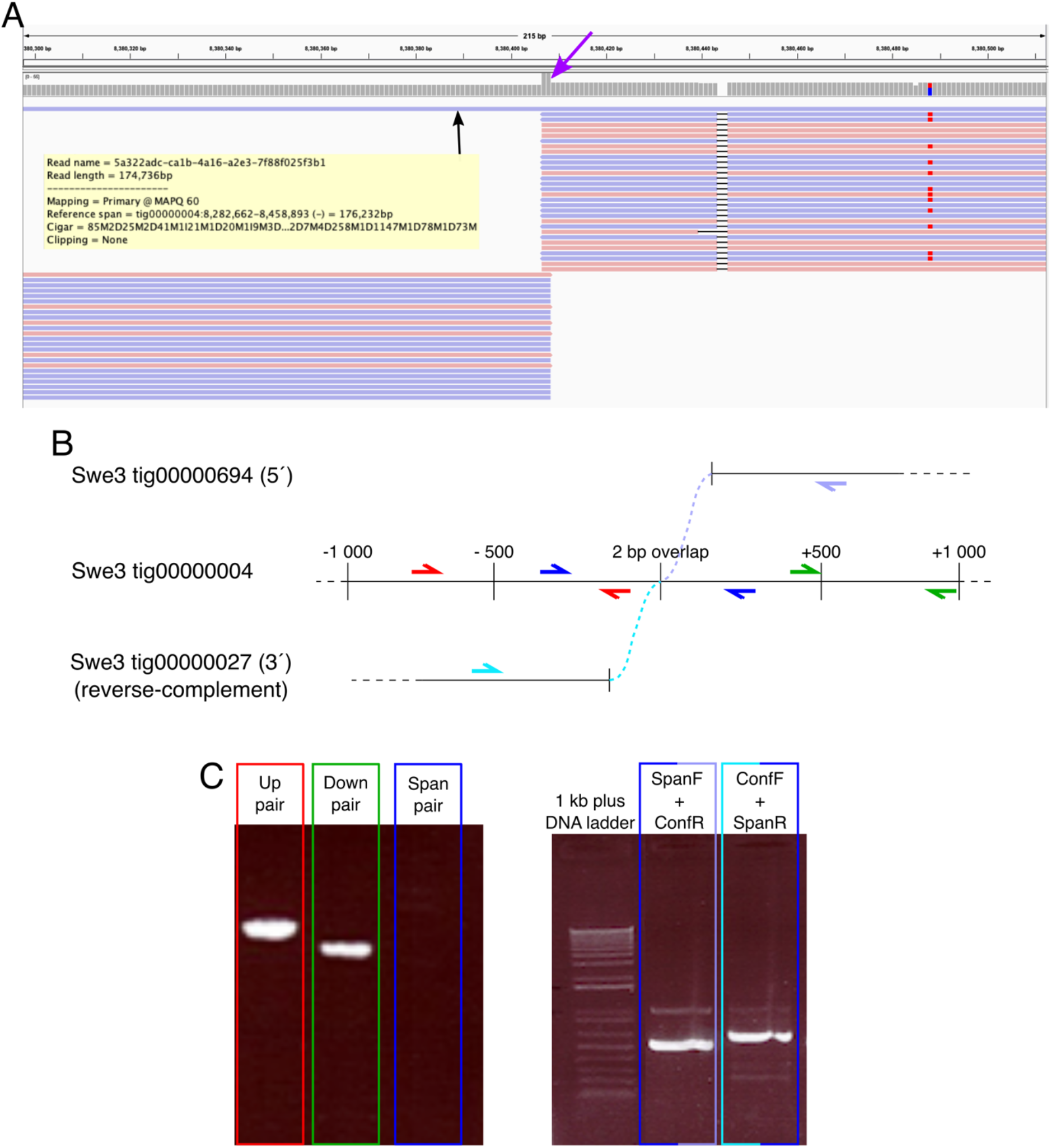
Detection of an error in the Swe3 initial assembly introduced by a long chimeric read. A. Mapping of the Swe3 reads, corrected and trimmed by Canu, on the Swe3 Canu assembly. The putative 175 kb chimeric read was identified by a break of synteny (not shown) and because of a sharp (ca. 2-fold) increase in coverage, spanning only 2 nucleotides, (purple arrow) on the contig tig0000004. B. Conceptual design of the PCR primers used to verify the assembly. Three pairs of primers were designed on the tig0000004: the Up pair (red), the Down pair (green) and the Span pair (blue). Two other primers were designed on the basis of the corrected assembly sequence (dotted coloured lines): SpanF in turquoise on the contig tig00000027 and SpanR in light purple on the contig tig00000695. C. PCR result of the different pairs used. Amplicons had the expected sizes.

**Supplementary Figure 4.**
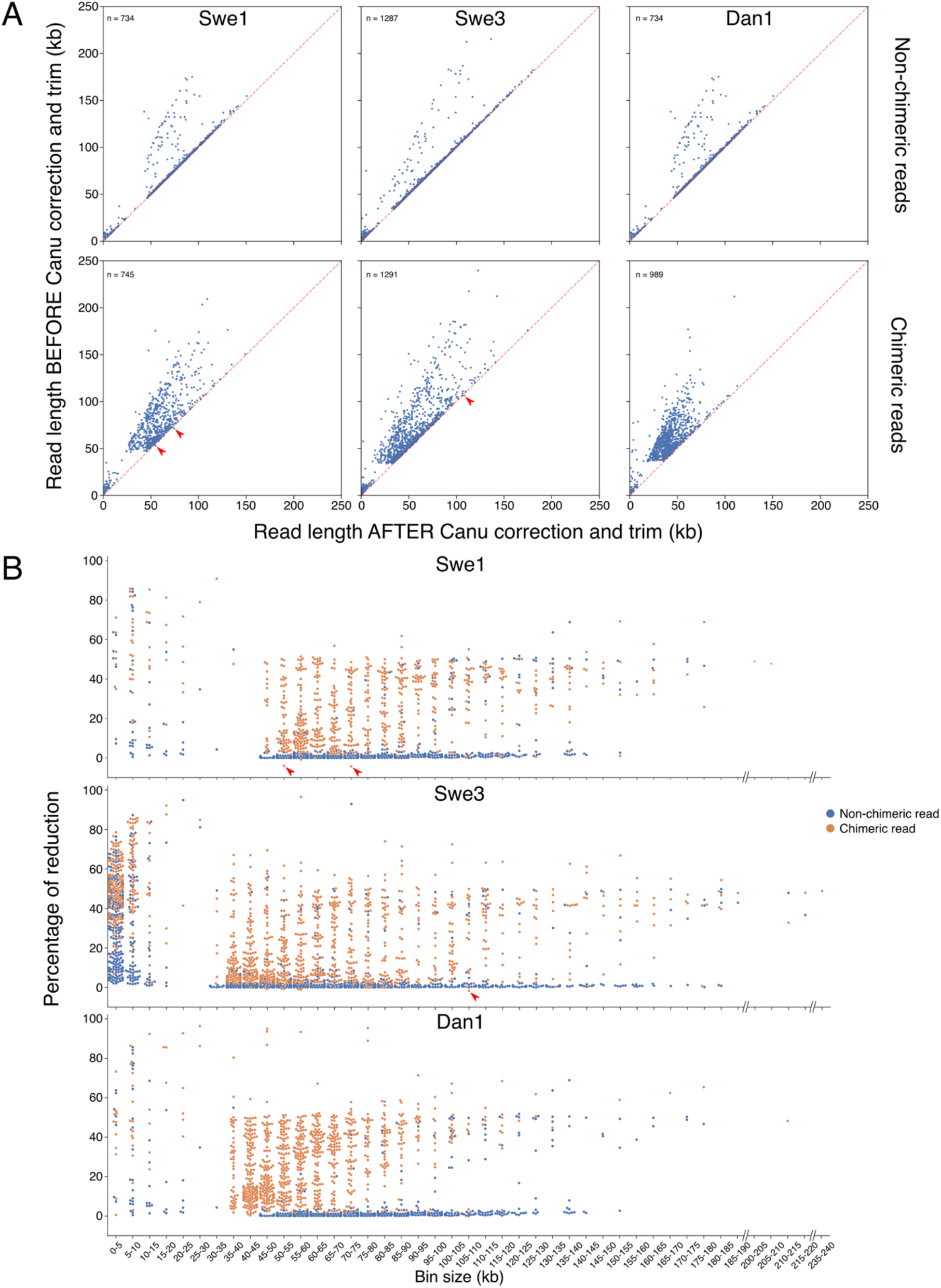
Canu correction of non-chimeric and chimeric reads. A. Scatter plots of the size of reads in the initial dataset (y-axis) and after Canu correction (x-axis) for non-chimeric (top) and chimeric (bottom) reads. Subsets of non-chimeric reads of a similar number (as indicated) and having the equivalent size distribution (before Canu correction, +/− 1% in length) as the chimeric read were used to allow a valid comparison. B. Percentage of read length reduction after Canu correction, in 5 kb bins. The bin size corresponds to the read size in the raw dataset. The red arrows highlight the rare chimeric reads that are longer after the correction step.

**Supplementary Figure 5.**
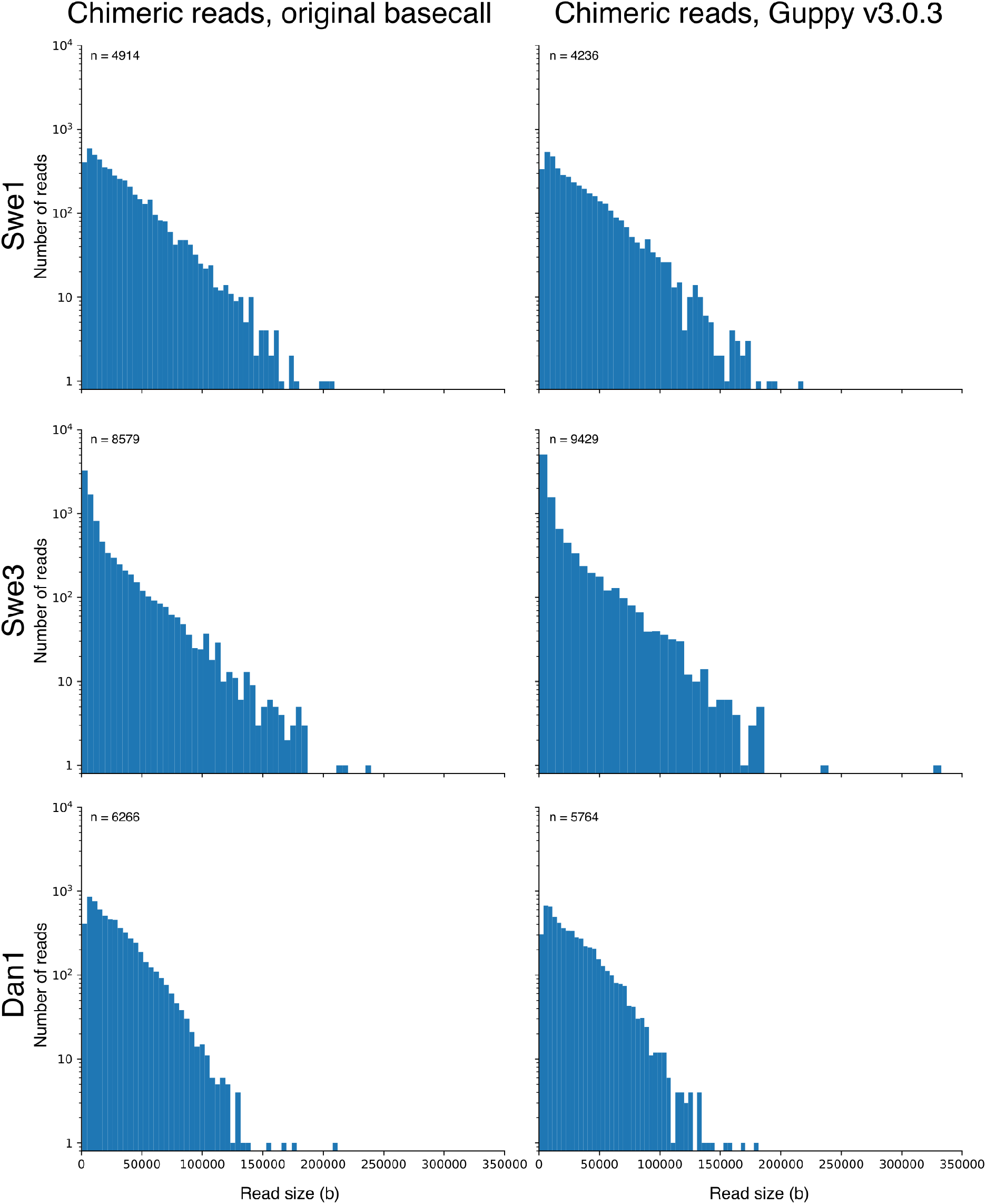
Size distribution of chimeric reads. Distributions in 5 kb bins of the size of the set of reads identified as chimeric by YACRD (see Methods), in the original dataset (left panels) and amongst reads basecalled by Guppy v3.0.3 (right panels).

**Supplementary Figure 6.**
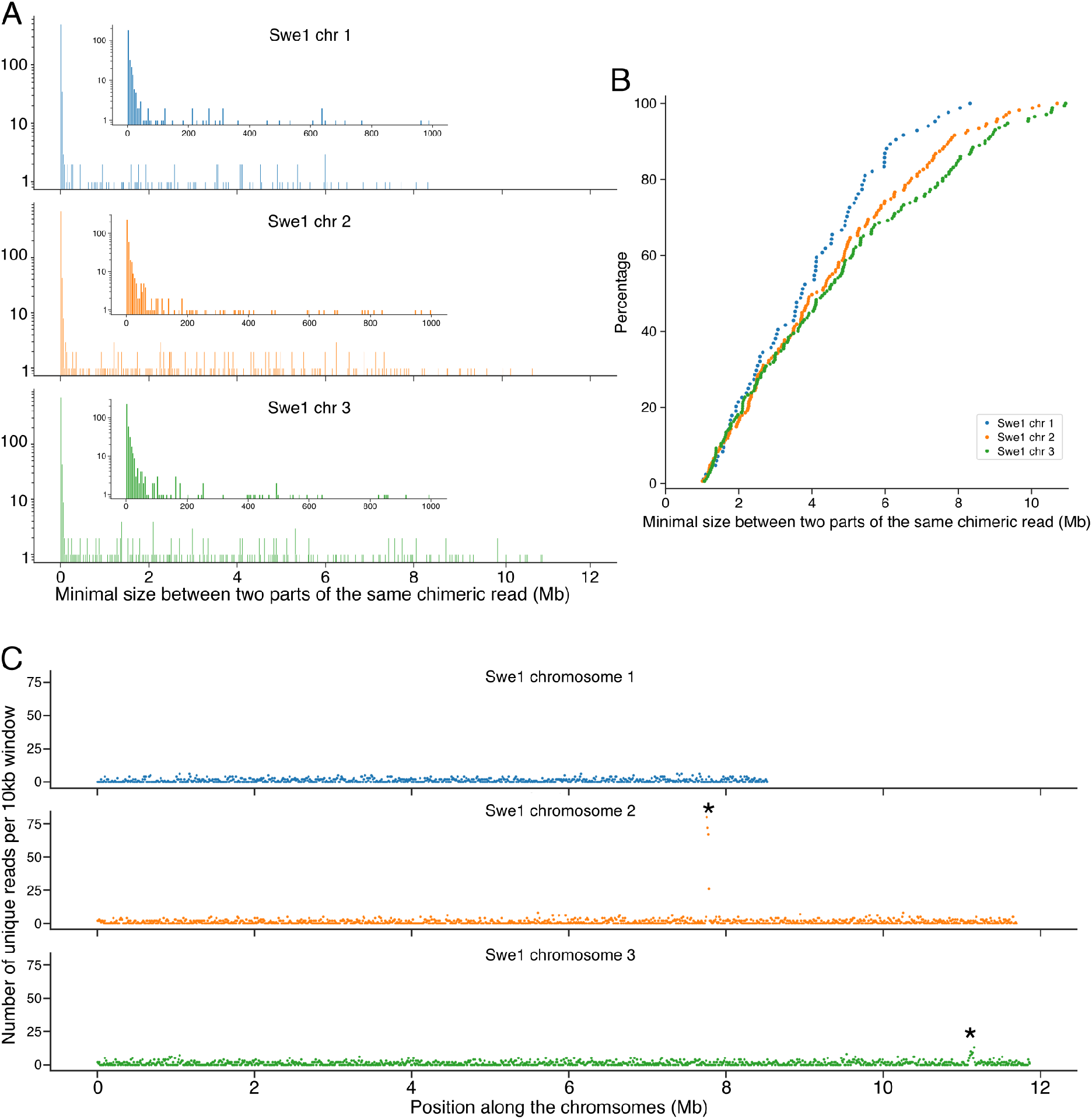
Analysis of the Swe1 chimeric reads. For each chimeric read, the 500 bp of sequence on each side of the putative break point was mapped onto the final Swe1 genome. A, Distribution in 500 equal bins of the interval between the two parts of each chimeric read. The inserts highlight reads that map within 1 kb of each other (in 200 bins). B, Cumulative distribution of the interval between the two parts of each chimeric read (for those separated by > 1 Mb). The distance separating the two parts is broadly spread. C. Distribution of mapped sequences from each side of the putative breakpoint for each chimeric read that mapped to a single chromosome. The peak on chromosome 2 corresponds to the site where the nuclear genome matches the mitochondrial DNA, and that on chromosome 3 to a highly repeated region containing tandem copies of rDNA. In both cases, these therefore reflect erroneous attribution of chimerism. There are thus no true hot spots for chimeric read breakpoints on any chromosome.

**Supplementary Figure 7.**
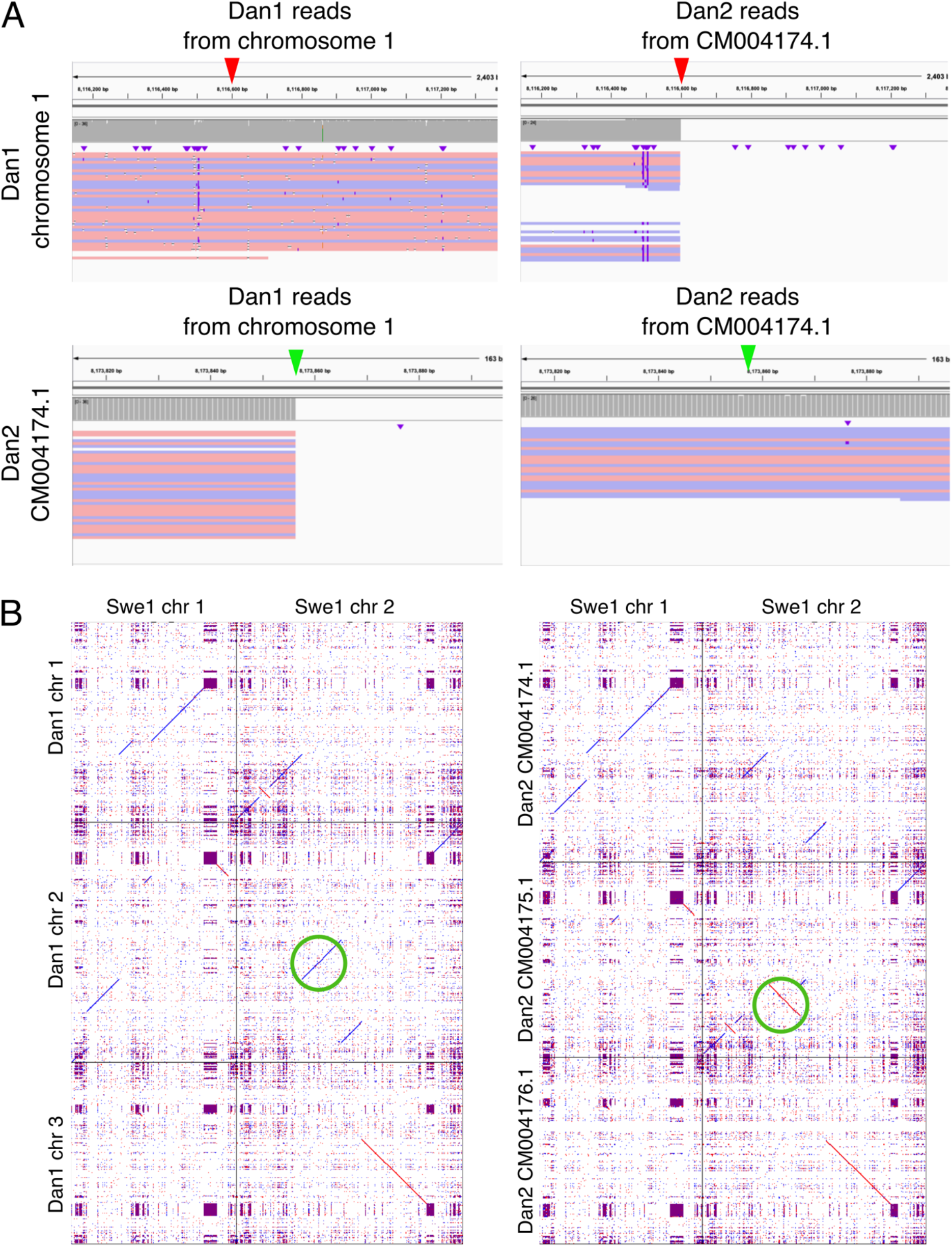
Identification of intra- and inter-chromosomal rearrangements between Dan1 and Dan2. A. Mapping of long reads from Dan1 (left panels) and Dan2 (right panels) on Dan1 chromosome 1 (top panels) and Dan2 chromosome 1 (CM004174.1; bottom panels). The arrowheads highlight points of discontinuity in the read coverage, consistent with chromosomal rearrangements between Dan1 and Dan2. B. Alignment of selected chromosomes of the final assemblies between Dan1 and Swe1 (left) and Dan2 and Swe1 (right). The unique difference in the orientation of part of Swe1 chromosome 2 between Dan1 and Dan2 is highlighted by the green circles.

**Supplementary Figure 8.**
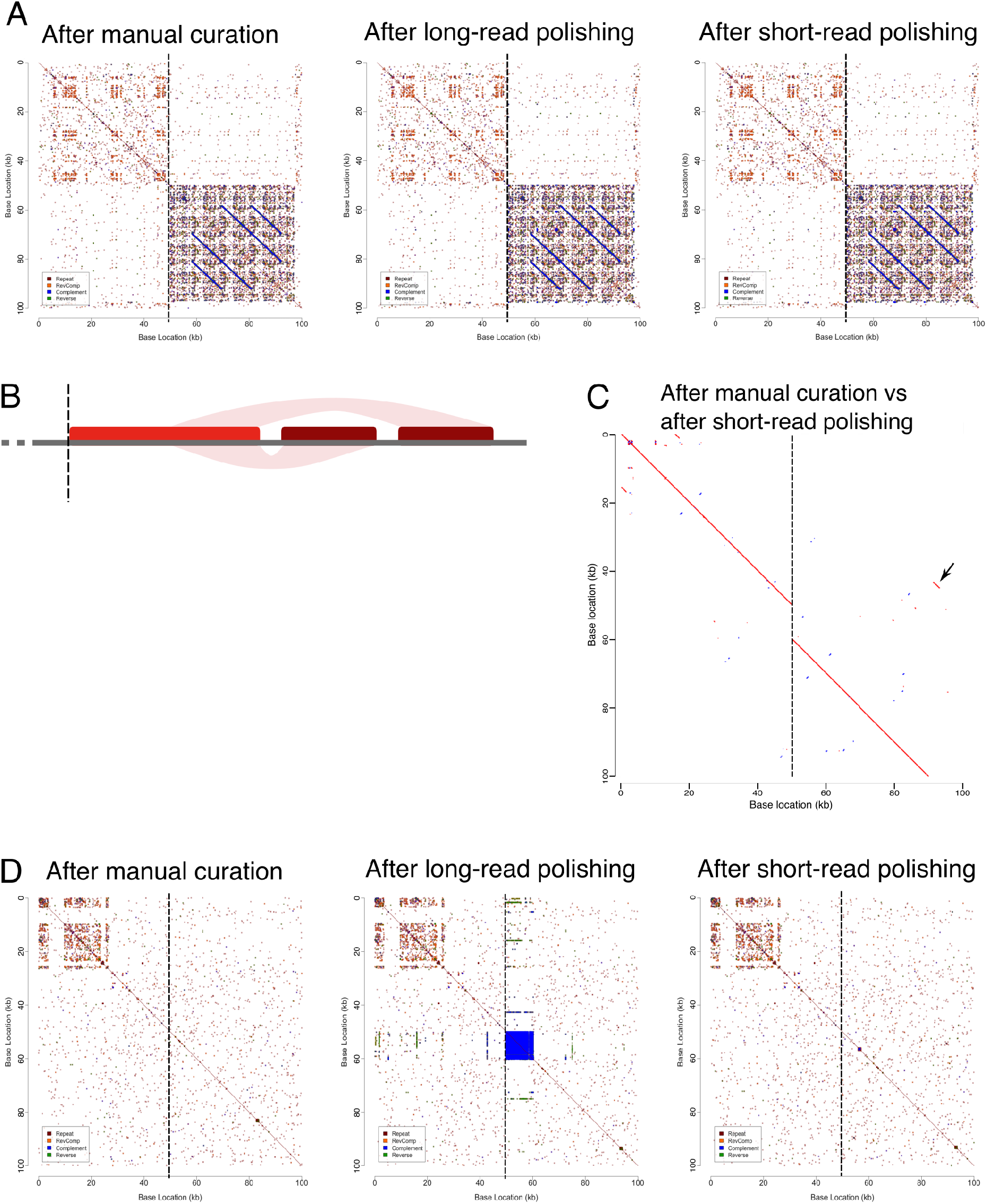
Analysis of the pattern of repeated sequences in the neighbourhood of polishing error in the Swe3 genome. A. Dot-plot based on k-mer identity (k = 11; See Supplementary Methods) of the neighbourhood of the *numts vs* itself, from the left to the right, before any polishing, after long-read polishing and after short-read polishing. The vertical dashed line at the centre of each plot indicates the start of the *numts*. B. A schematic representation of the large-scale genomic organization in the same region. The rectangles (red and dark red) represent *numts* (identified by BLASTN against the Swe3 mitochondrial genome) and the ribbons show the regions of sequence similarity (>99.5%) between them. For the Swe1 assembly, one dark-red rectangle is absent, while in the Swe2 assembly, the two dark-red rectangles are absent. C. An alignment of the area before any polishing (x-axis) and after short-read polishing (y-axis) reveals the re-inclusion of this missing sequence in the final assembly. The arrow highlights a 1.2 kb region that is duplicated on both sides of the sequence discontinuity (vertical dashed line). D. Dot-plots based on k-mers (k = 11) of the same region. The blue square reflects the presence of a region of low-complexity repeated sequences. The pattern at the top left hand of the plots reflect a region of low-complexity sequence that is not shared across the sequence discontinuity (vertical dashed line).

**Supplementary table 1:** Coverage, genome length and results of the tBLASTn

|  | Set of reads | Swe1 | Swe3 | Dan1 |
| --- | --- | --- | --- | --- |
| Coverage (fold) | Long read - Guppy 1.5.1 | 136 | 94 | 150 |
|  | Long read - Canu corrected | 36 | 36 | 36 |
|  | Long read - Guppy 3.0.3 | 140 | 97 | 155 |
|  | Short read | 406 | 430 | 482 |
| Genome length (bp) | Canu after curation | 31971578 | 32076624 | 31919747 |
|  | Long-read polishing | 32121045 | 32271010 | 32120031 |
|  | Short-read polishing | 32117370 | 32271155 | 32095731 |
| TBLASTN result for the 305 benchmarking genes | Canu after curation | 43 (14%) | 48 (16%) | 42 (14%) |
|  | Long-read polishing | 236 (77%) | 248 (81%) | 152 (50%) |
|  | Short-read polishing | 305 (100%) | 305 (100%) | 305 (100%) |

The coverage for each set of reads was computed as the mean depth for the assemblies after short-read polishing, using Samtools coverage v1.10.

**Supplementary Table 2:**
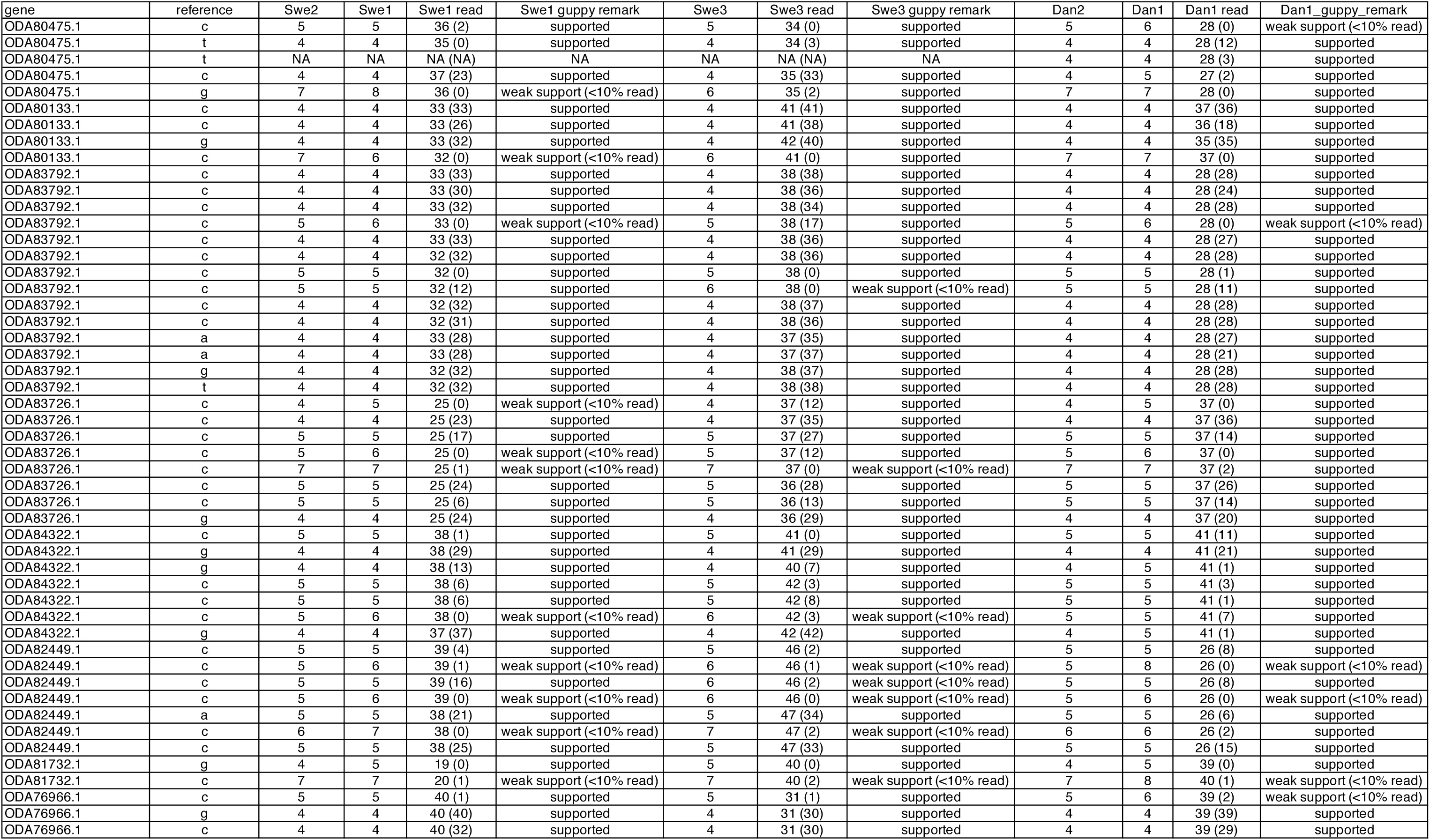
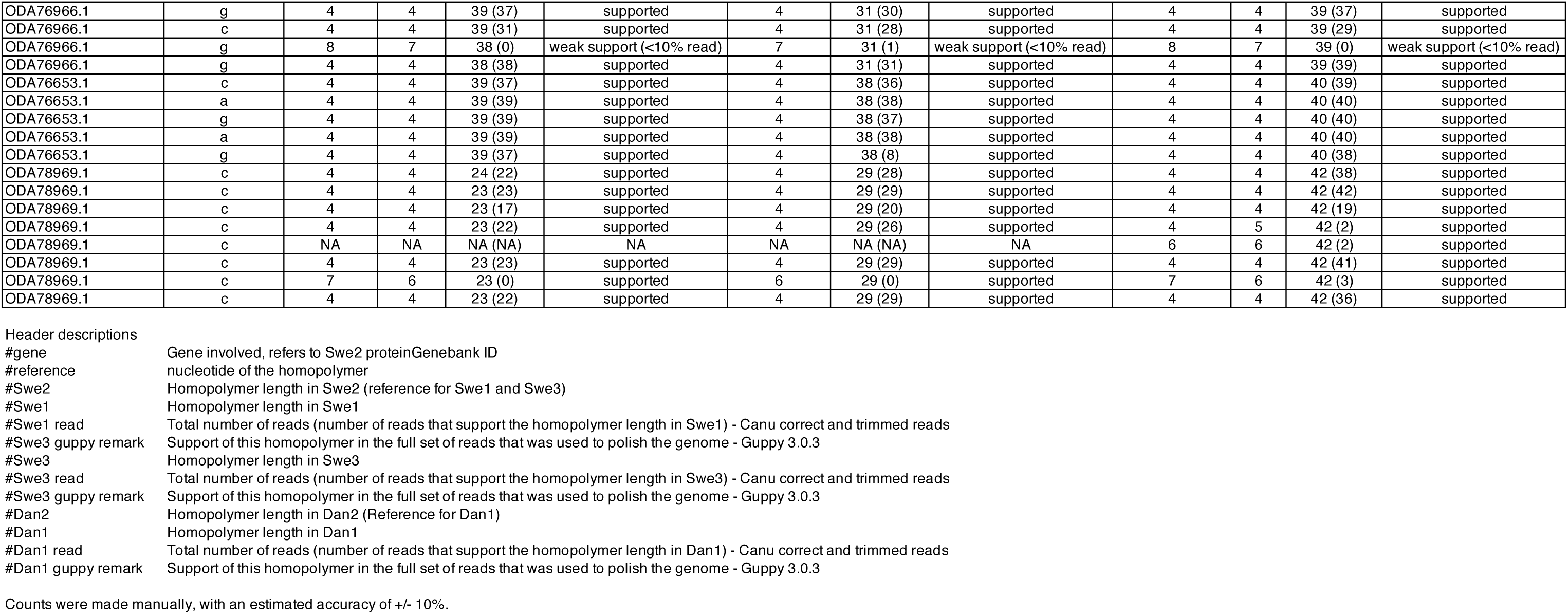
Examples of read support for different homopolymer sequences in 10 representative genes.

## Supplementary methods

### GenBank query

We queried the Assembly database on the NCBI website with: (“minion”[Sequencing Technology] OR “nanopore”[Sequencing Technology] OR “nanopore minion”[Sequencing Technology] OR “nanopore technologies”[Sequencing Technology] OR “nanopore technology”[Sequencing Technology] OR “nanpore”[Sequencing Technology] OR “ont”[Sequencing Technology] OR “ont minion”[Sequencing Technology] OR “oxford”[Sequencing Technology] OR “oxford nanopore”[Sequencing Technology] OR “oxford nanopore minion”[Sequencing Technology] OR “oxford nanopore technologies”[Sequencing Technology] OR “oxford nanopore technology”[Sequencing Technology]) NOT (“illumina” OR “bgi” OR “pacbio” OR “pacbio rs” OR “pacbio rs ii” OR “pacbio rsii” OR “pacbio sequel” OR “pacbio smrt” OR “pacific biosciences” OR “sequel” OR “hybrid” OR “hybrid assembly”) AND (latest[filter] OR “latest genbank”[filter]) AND (all[filter] NOT “derived from surveillance project”[filter]) AND (all[filter] NOT anomalous[filter]) The accuracy of this query depends on the assembly metadata in GenBank.

### K-mer based dot-plots

The dot-plots based on k-mers were computed with the script *repaver.r* (available at https://gitlab.com/gringer/bioinfscripts) The version used here was the commit 0854f3af76cb5bc5a7e0a2b4a032518f51e210fa, with the parameters *-k 11 -style dotplot*.

### Search for Dan1 / Dan2 transposable element at the breakpoints

A first analysis was conducted with the Dfam (https://dfam.org/home) to search for transposable elements (TE). The parameter *organism* was set to *other*. No TE was present in the neighbourhood of the different breakpoints. We also mined the two genomes for TE with transposonPSI (http://transposonpsi.sourceforge.net/) with default parameters. Here also, no TE was present within the 50kb surrounding each breakpoint.

### PCR

PCR was performed to test the putative chimeric nature of a Canu contig in the Swe3 assembly, with primers UpF (AACTGTGTCTAACTAGCCCG), UpR (AGGGTCCTCATAAACTTGGC), DownF (TGTATCAGGTTCCCGAATGG), DownR (CTAGGCTGGGGAATCTTCTG), SpanF (CCATCAACTTCAGCTGCTC), SpanR (CTCCTCAATCTCCCTCTCGG), ConfF (ATCGGCGACTACCTGCAC), ConfR (CGTTCCATCGTTACCACAGC). PCR reactions were run according to the GoTaq® G2 Flexi DNA polymerase instructions (Promega), with 50 ng of template DNA, 1 mM of each forward and reverse primers, in a final volume of 25 *μ*L. The reaction started by initial denaturation at 95°C for 2 min, followed by 30 amplification cycles (95°C for 30 sec, 60°C for 30 sec and 72°C for 30 sec), and a final elongation for 5 min at 72°C.

## References

1. Jansson HB, Jeyaprakash A, Zuckerman BM. Differential Adhesion and Infection of Nematodes by the Endoparasitic Fungus *Meria coniospora* (Deuteromycetes). Appl Environ Microbiol. 1985;49:552–5.

2. Pujol N, Link EM, Liu LX, Kurz CL, Alloing G, Tan MW, et al. A reverse genetic analysis of components of the Toll signalling pathway in *Caenorhabditis elegans*. Curr Biol. 2001;11:809–21.

3. Dierking K, Polanowska J, Omi S, Engelmann I, Gut M, Lembo F, et al. Unusual regulation of a STAT protein by an SLC6 family transporter in *C. elegans* epidermal innate immunity. Cell Host Microbe. 2011;9:425–35.

4. Labed SA, Omi S, Gut M, Ewbank JJ, Pujol N. The pseudokinase NIPI-4 is a novel regulator of antimicrobial peptide gene expression. PloS One. 2012;7:e33887.

5. Lebrigand K, He LD, Thakur N, Arguel M-J, Polanowska J, Henrissat B, et al. Comparative Genomic Analysis of *Drechmeria coniospora* Reveals Core and Specific Genetic Requirements for Fungal Endoparasitism of Nematodes. PLoS Genet. 2016;12:e1006017.

6. Zhang L, Zhou Z, Guo Q, Fokkens L, Miskei M, Pócsi I, et al. Insights into Adaptations to a Near-Obligate Nematode Endoparasitic Lifestyle from the Finished Genome of *Drechmeria coniospora*. Sci Rep. 2016;6:23122.

7. Canu 1.8 documentation. https://canu.readthedocs.io/en/latest/faq.html#my-circular-element-is-duplicated-has-overlap. Accessed 15 Nov 2019

8. Kolmogorov M. Fast and accurate de novo assembler for single molecule sequencing reads: fenderglass/Flye. https://github.com/fenderglass/Flye. Accessed 3 June 2019

9. Schechtman MG. Characterization of telomere DNA from *Neurospora crassa*. Gene. 1990;88:159–65.

10. Hazkani-Covo E, Zeller RM, Martin W. Molecular Poltergeists: Mitochondrial DNA Copies (numts) in Sequenced Nuclear Genomes. PLoS Genet. 2010;6:e1000834.

11. Argueso JL, Westmoreland J, Mieczkowski PA, Gawel M, Petes TD, Resnick MA. Double-strand breaks associated with repetitive DNA can reshape the genome. Proc Natl Acad Sci U S A. 2008;105:11845–50.

12. Sun S, Yadav V, Billmyre RB, Cuomo CA, Nowrousian M, Wang L, et al. Fungal genome and mating system transitions facilitated by chromosomal translocations involving intercentromeric recombination. PLoS Biol. 2017;15:e2002527.

13. TransposonPSI: An Application of PSI-Blast to Mine (Retro-)Transposon ORF Homologies. http://transposonpsi.sourceforge.net. Accessed 1 Apr 2020

14. Hubley R, Finn RD, Clements J, Eddy SR, Jones TA, Bao W, et al. The Dfam database of repetitive DNA families. Nucleic Acids Res. Oxford Academic; 2016;44:D81–9.

15. Senol Cali D, Kim JS, Ghose S, Alkan C, Mutlu O. Nanopore sequencing technology and tools for genome assembly: computational analysis of the current state, bottlenecks and future directions. Brief Bioinform. 2018;20:1542–59.

16. Scheunert A, Dorfner M, Lingl T, Oberprieler C. Can we use it? On the utility of de novo and reference-based assembly of Nanopore data for plant plastome sequencing. PloS One. 2020;15:e0226234.

17. White R, Pellefigues C, Ronchese F, Lamiable O, Eccles D. Investigation of chimeric reads using the MinION. F1000Res. 2017;6:631.

18. Eccles D, Chandler J, Camberis M, Henrissat B, Koren S, Le Gros G, et al. De novo assembly of the complex genome of *Nippostrongylus brasiliensis* using MinION long reads. BMC Biol. 2018;16:6.

19. Schmid M, Frei D, Patrignani A, Schlapbach R, Frey JE, Remus-Emsermann MNP, et al. Pushing the limits of de novo genome assembly for complex prokaryotic genomes harboring very long, near identical repeats. Nucleic Acids Res. 2018;46:8953–65.

20. Watson M, Warr A. Errors in long-read assemblies can critically affect protein prediction. Nature Biotechnology. 2019;37:124–6.

21. Wick RR, Judd LM, Holt KE. Performance of neural network basecalling tools for Oxford Nanopore sequencing. Genome Biol. 2019;20:129.

22. Somerville V, Lutz S, Schmid M, Frei D, Moser A, Irmler S, et al. Long-read based de novo assembly of low-complexity metagenome samples results in finished genomes and reveals insights into strain diversity and an active phage system. BMC Microbiol. 2019;19:143.

23. Dal Molin A, Minio A, Griggio F, Delledonne M, Infantino A, Aragona M. The genome assembly of the fungal pathogen *Pyrenochaeta lycopersici* from Single-Molecule Real-Time sequencing sheds new light on its biological complexity. PLoS One. 2018;13:e0200217

24. Jain M, Koren S, Miga KH, Quick J, Rand AC, Sasani TA, et al. Nanopore sequencing and assembly of a human genome with ultra-long reads. Nat Biotechnol. 2018;36:338–45.

25. Simao FA, Waterhouse RM, Ioannidis P, Kriventseva EV, Zdobnov EM. BUSCO: assessing genome assembly and annotation completeness with single-copy orthologs. Bioinformatics. 2015;31:3210–2.

26. He LD, Ewbank JJ. Polyethylene Glycol-mediated Transformation of *Drechmeria coniospora*. Bio-Protoc. 2017;7:e2157.

27. Fungal DNA extraction protocol. https://www.pnas.org/content/pnas/suppl/2018/01/08/1715954115.DCSupplemental/pnas.1715954115.sapp.pdf. Accessed 16 Apr 2020.

28. Kjærbølling I, Vesth TC, Frisvad JC, Nybo JL, Theobald S, Kuo A, et al. Linking secondary metabolites to gene clusters through genome sequencing of six diverse *Aspergillus* species. Proc Natl Acad Sci. 2018;115:E753–61.

29. Quick J. Ultra-long read sequencing protocol for RAD004 v3. https://www.protocols.io/view/ultra-long-read-sequencing-protocol-for-rad004-mrxc57n. Accessed 25 Oct 2019

30. Wick R. Porechop. https://github.com/rrwick/Porechop. Accessed 3 Dec 2019

31. Marijon P, Chikhi R, Varré J-S. yacrd and fpa: upstream tools for long-read genome assembly. bioRxiv. 2019;674036.

32. Kiełbasa SM, Wan R, Sato K, Horton P, Frith MC. Adaptive seeds tame genomic sequence comparison. Genome Res. 2011;21:487–93.

33. Krzywinski M, Schein J, Birol İ, Connors J, Gascoyne R, Horsman D, et al. Circos: An information aesthetic for comparative genomics. Genome Res. 2009;19:1639–45.

34. Li H. Minimap2: pairwise alignment for nucleotide sequences. Bioinformatics. 2018;34:3094–100.

35. Li H, Handsaker B, Wysoker A, Fennell T, Ruan J, Homer N, et al. The Sequence Alignment/Map format and SAMtools. Bioinforma Oxf Engl. 2009;25:2078–9.

36. Thorvaldsdóttir H, Robinson JT, Mesirov JP. Integrative Genomics Viewer (IGV): high-performance genomics data visualization and exploration. Brief Bioinform. 2013;14:178–92.

37. Koren S, Walenz BP, Berlin K, Miller JR, Bergman NH, Phillippy AM. Canu: scalable and accurate long-read assembly via adaptive k-mer weighting and repeat separation. Genome Res. 2017;27:722–36.

38. Altschul SF, Gish W, Miller W, Myers EW, Lipman DJ. Basic local alignment search tool. J Mol Biol. 1990;215:403–10.

39. Katoh K, Standley DM. MAFFT multiple sequence alignment software version 7: improvements in performance and usability. Mol Biol Evol. 2013;30:772–80.

40. Gouy M, Guindon S, Gascuel O. SeaView Version 4: A Multiplatform Graphical User Interface for Sequence Alignment and Phylogenetic Tree Building. Mol Biol Evol. 2010;27:221–4.

41. Vaser R, Sović I, Nagarajan N, Šikić M. Fast and accurate de novo genome assembly from long uncorrected reads. Genome Res. 2017;27:737–46.

42. Kurtz S, Phillippy A, Delcher AL, Smoot M, Shumway M, Antonescu C, et al. Versatile and open software for comparing large genomes. Genome Biol. 2004;5:R12.

43. Bolger AM, Lohse M, Usadel B. Trimmomatic: a flexible trimmer for Illumina sequence data. Bioinformatics. Oxford Academic; 2014;30:2114–20.

44. Li H. Aligning sequence reads, clone sequences and assembly contigs with BWA-MEM. ArXiv 2013;1303.3997

45. Walker BJ, Abeel T, Shea T, Priest M, Abouelliel A, Sakthikumar S, et al. Pilon: An Integrated Tool for Comprehensive Microbial Variant Detection and Genome Assembly Improvement. PLoS One. 2014;9:e112963.

